# Germline sequence variation within the ribosomal DNA is associated with human complex traits

**DOI:** 10.1101/2025.02.06.635840

**Authors:** Francisco Rodriguez-Algarra, Maia Cooper, Faraz K Mardakheh, David M Evans, Vardhman K Rakyan

## Abstract

The ribosome is one of the core macromolecules in the cell. The ribosomal RNAs (rRNA), which are essential components of the ribosome, are coded by the multi-copy ribosomal DNA (rDNA). Despite its highly conserved function, the rDNA displays substantial variation within all species analysed to date. This variation comprises both inter-individual differences in total copy number (CN) as well as inter- and intragenomic sequence variation in the form of single nucleotide variants (SNV) and insertions/deletions (INDELs) across rDNA copies. Whether germline variation of rDNA sequence associates with phenotypic traits in humans is, to date, unknown. Here, using the UK Biobank whole genome sequencing data, we first derive a high confidence list of rDNA-associated SNVs and INDELs that we validate in multiple ways. Using this list, we show that specific rDNA variants associate with several human traits. In particular, traits associated with body size appear enriched in variants within the Expansion Segment 15L region in the 28S rRNA. The strength of these associations does not diminish when accounting for the total rDNA CN of each individual. Our work represents the first large-scale association analysis of human traits with germline sequence variation in the rDNA, a source of human complex trait-relevant genetic variation that has thus far been largely ignored.

## Introduction

The ribosome is the cell’s translational apparatus and therefore one of the core macromolecules of life (**Fig. 1A**). In humans, the mature ribosome is comprised of ∼80 different proteins, and 4 different ribosomal RNAs (rRNA) – the 5S, 18S, 5.8S, and 28S – which are the most abundant RNA species in the cell. The high levels of rRNA expression are supported by the multi-copy nature of the ribosomal DNA. Interestingly, because of elevated homologous recombination amongst different rDNA clusters, rDNA copy number (CN) displays significant inter-individual variation (∼200 – 600 copies within humans)^[1,2]^. Furthermore, the different copies of rDNA are genetically variable, containing single nucleotide variants (SNVs) and insertion/deletions (INDELs)^[1,3,4]^, demonstrating that concerted evolution only partially homogenises the sequences across different rDNA copies^[5]^.

**Figure 1A.**
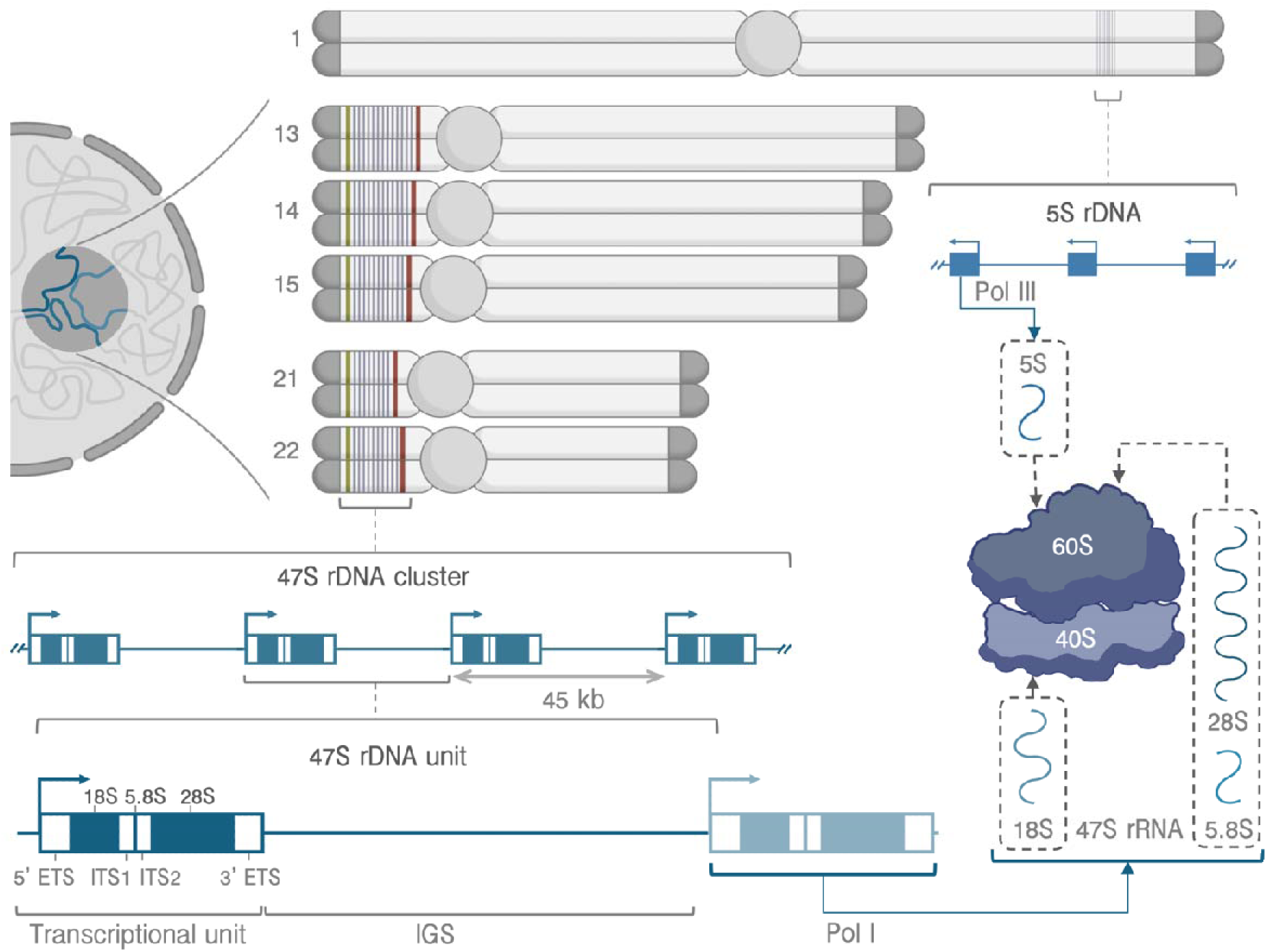
Schematic representation of the human rDNA loci. Adapted from [7].

But what is the phenotypic impact of naturally occurring genetic variation within human rDNA? Until recently, this was difficult to answer because of the genetic complexity of rDNA clusters, and because rDNA is either not represented on commercial microarray platforms, or typically excluded in standard computational pipelines for analysing sequencing data. This is now changing with increasing availability of large-scale whole genome sequencing (WGS) datasets, and development of novel computational approaches^[4,6]^.

Phenotypic associations of human inter-individual rDNA genetic variation have been explored in the context of both somatic and germline variation^[7]^. Somatic genetic variation of human rDNA has been noted in several biological contexts, most robustly in cancer^[8,9]^, and also during aging^[10]^, and in response to environmental insults^[11,12]^, albeit in very small scale studies. Somatic variation includes both copy number alterations and, in cancer, apparent differential expression of specific variant copies of the rDNA^[4]^. However, in all these cases, rDNA genetic variation is most likely a downstream event, and it remains unclear whether somatic genetic variation of rDNA, including that which arises spontaneously, has any impact on trait outcomes.

On the other hand, trait-associated germline variation must either directly or indirectly, influence the trait or disease in question. For germline rDNA genetic variation, we recently reported the first robust evidence for a genetic association between germline rDNA CN variation and a variety of human complex traits, including neutrophil counts and kidney disease in the UK Biobank (UKB)^[6]^, and body mass in a separate cohort^[13]^. Importantly, we also established that rDNA CN variation is unlikely to be influenced by common genetic variation elsewhere in the genome.

However, rDNA copies are not genetically identical, harbouring both intra- and inter-individual SNVs and INDELs (hereby collectively referred to as ‘variants’). Previous studies have found that these variants are located throughout the rDNA unit. This includes within the rRNA subunits that are incorporated into the mature ribosome, leading to the intriguing possibility of rRNA-based ribosomal hetereogeneity and consequently impacts on translational outcomes^[14–16]^, as has already been reported for bacteria^[17,18]^.

The recent availability of WGS data from large, publicly accessible human biobank studies has offered unprecedented opportunities to perform discovery analyses of the association between rDNA genetic variation and human traits. Here, we report an in-depth trait-association analysis of rDNA genetic variants in the UKB WGS data^[19–21]^. We first derive a high confidence list of rDNA-associated SNVs and INDELs that we validate in multiple ways. Using this list, we show that rDNA variants are associated with several human traits, most notably with body size-related phenotypes. Our work constitutes the first large-scale association analysis of human traits germline sequence variation in the rDNA, which represents a source of human complex trait-relevant germline genetic variation that has thus far been ignored.

## Results

### Selection of rDNA sequence variants for association analyses in UKB

Due to the hundreds of rDNA units that exist in the human genome, SNVs and INDELs in the rDNA may be present in proportions other than 100:0, 50:50, or 0:100 like variants in the single copy regions of the genome. Rather, each rDNA sequence variant may exist, within an individual, as any relative proportion from 0% to 100% depending on how many of all rDNA units harbour it (**Fig. 1B**). For instance, consider variant 7980:G>A where virtually every individual harbours rDNA units with Gs and As at that position. In particular, if one individual with a total rDNA CN of 400 had 300 of these containing a G (the reference allele) and the rest an A, we would record the variation of that individual at 7980:G>A as an intragenomic variant frequency of 0.25 (thus 25%). That value is the one that would be then used in further analyses. This property of rDNA variation impedes the use of conventional variant calling tools, which has in turn likely contributed to the wide discrepancies between the sets of variants reported in previous studies^[1,3,4]^. Since our main goal is to determine whether any rDNA sequence variation within the 47S transcriptional unit associates with human phenotypes and not necessarily determine how many variant positions exist in the region, we have devoted substantial efforts to establish a set of reliable rDNA variants in the UKB cohort.

**Figure 1B.**
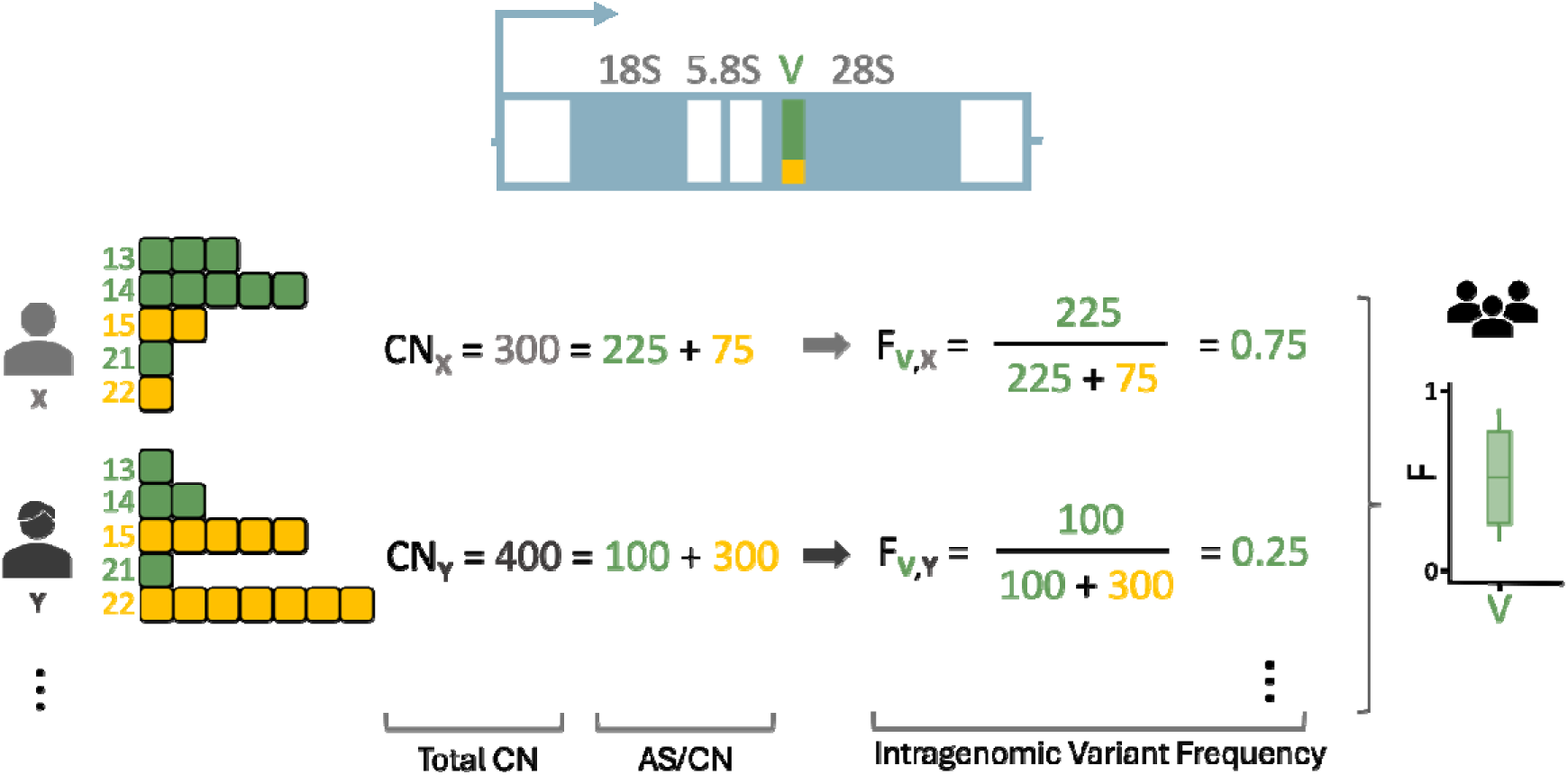
Toy example of sequence variation in the rDNA. Given a variant position V within the 28S with two potential alleles G (yellow, reference) and A (green), and assuming for simplicity that the alleles at this position are chromosome-specific, an individual X might harbour 225 of their total CN_X_ = 300 rDNA units with an A at that position (each square representing 25 rDNA units). In that case, their allele-specific copy number for the variant V:G>A is AS/CN_V,X_ = 225 and their corresponding intragenomic variant frequency is thus F_V,X_ = 0.75. Another individual Y, on the other hand, might only harbour an A in 100 of their total CN_Y_ = 400 rDNA unit. Their allele-specific copy number is hence AS/CN_V,Y_ = 100 and their corresponding intragenomic variant frequency is F_V,Y_ = 0.25. Across the population, the intragenomic variant frequencies at that position will follow a continuous distribution.

#### Monozygotic twin pairs as testbed for the UKB rDNA variation calling pipeline

The sheer scale of the UKB cohort adds a further layer of complexity to the analysis. Feasibly analysing hundreds of thousands of WGS samples requires some compromises, both in which subset of reads from the libraries we process and how this processing occurs, which may affect the reliability of the variant calls. On the other hand, the size of the dataset also offers an excellent opportunity to validate our methodological choices given the multiple pairs of monozygotic (MZ) twins included in the cohort. These display a very high correlation in their total number of rDNA copies^[6]^. Since sequencing centre has a strong impact on the total number of rDNA CN estimated for each UKB participant^[6]^, we consider all 49 white British (WB) MZ twin pairs with both individuals sequenced in the same centre and release. We then use those to validate the reference assembly for realignment, the quality of the variant calls themselves, and the effect of the extraction of rDNA analogue reads from the pre-existing alignments (**Fig. 1C**).

**Figure 1C.**
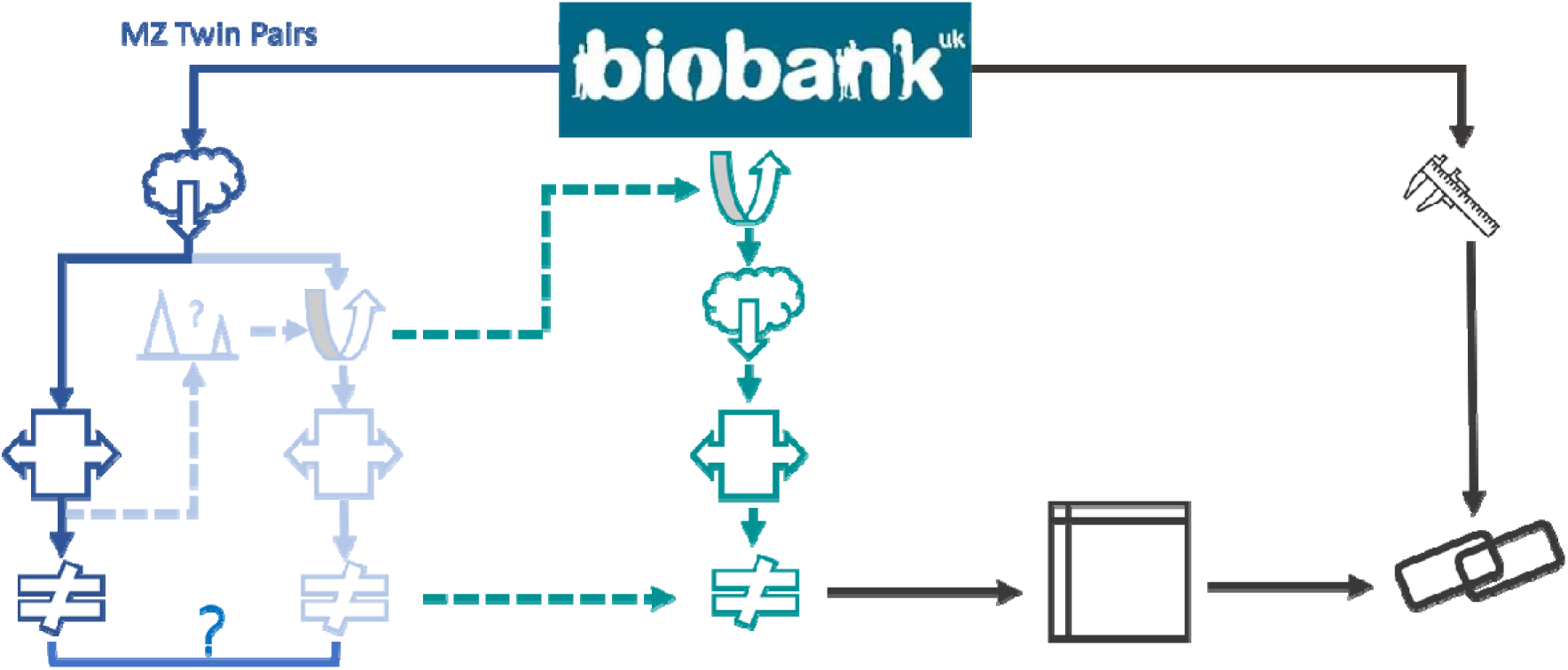
Schematic representation of the validation and analysis pipelines employed in this study. To select a reliable set of rDNA variants to analyse, alignment files for white British UKB participants in MZ twin pairs were retrieved in full. These were then processed in two different ways. First, the full alignment files were realigned to diverse reference sequences. The result of these realignments was used both to compare the resulting variant frequencies and to determine the location of rDNA analogue regions in Hg38. The reliability of the variant calls themselves was determined comparing the estimates within twin pairs. Second, reads from the pre-existing alignments mapping to the identified regions were extracted and realigned in turn to determine the reliability of the extraction procedure. These validation steps then informed the procedure employed in the full cohort, where rDNA analogue reads were extracted prior to alignment. The resulting variant calls were then tabulated and regressed against a range of UKB traits to determine putative phenotypic associations with intragenomic variant frequencies.

#### Employing an rDNA-only realignment reference only affects low-frequency variant calls

In the past, we and others have processed WGS data aligning it to a tailored reference assembly including both a single rDNA unit and the entire consensus sequence for the rest of the genome (e.g., Hg38)^[6,13,22–24]^. This was intended to reduce the likelihood of spurious alignments from reads arising from elsewhere in the genome affecting the rDNA results. Other research groups tackling rDNA variation, however, opted for aligning the libraries solely to an rDNA unit or rRNA gene sequence consensus^[1,3,4]^. This reduces the computational burden of the analysis, but at the risk of less reliable variation calls. Given the size of the UKB cohort, we wondered whether the increase in efficiency may outweigh the potential noise introduced in the calls. To this end, we realigned the full WGS data for the 98 UKB individuals in the considered WB MZ twin pairs to both the same tailored whole-genome plus rDNA reference we used in the past^[6,13,23]^ and the looped rDNA consensus sequence included in the tailored reference. As **Fig. 2A** shows, differences in the intragenomic variant frequencies from the rDNA transcriptional unit derived from these two methods only arise in variants within the lower range of frequencies (Pearson’s R = 0.99, p = 0). We observe similar results when employing other publicly-available whole-genome plus rDNA references instead^[24]^ (**fig. S1**). We thus decided to introduce a pragmatic variant filtering step, retaining only variants appearing in at least one twin pair in high-confidence calls with intragenomic frequency between 10 and 90%. 457 variants across the rDNA transcriptional unit satisfy this requirement.

**Figure 2A.**
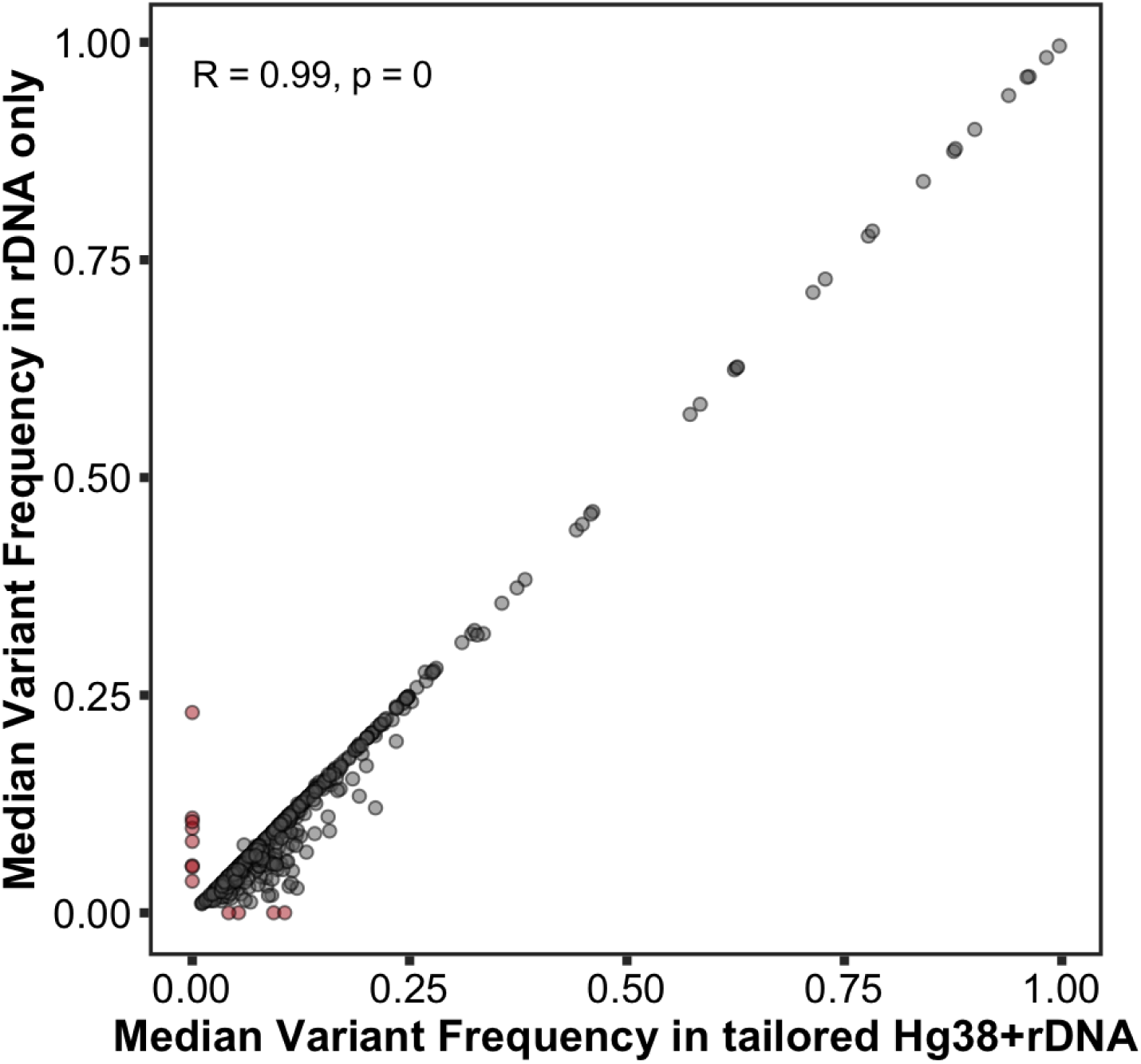
Comparison between the median frequency for each identified variant in the rDNA transcriptional unit estimated from realignments of UKB MZ twin samples to the full tailored Hg38+rDNA consensus assembly and a looped KY962518.1 rDNA reference. Red dots indicate variants that were not detected at any frequency in only one of the approaches.

#### Intra-twin-pair frequency comparisons reveal apparent sequencing artefacts

The main benefit of relying on MZ twin pairs for variant selection and validation is that they can act as replicates to test the reliability of the variant calls, since we expect intragenomic frequency estimates to match within each pair. To this end, for each variant satisfying the criteria defined above, we calculate the correlation between the intragenomic frequency estimates for the two individuals of each pair. For most variants, this comparison looks like the example in the left panel of **Fig. 2B**, with a Pearson’s correlation value of approximately 1, as we would expect. For other variants, however, the observed correlation is substantially lower, with virtually unrelated frequency estimates within each pair, such as in the example depicted in the right panel of **Fig. 2B**. Overall, approximately 16.8% of variants (77 out of 457) display intra-twin-pair frequency correlations below 0.8, with apparent clusters of diverging variants throughout the rDNA transcriptional unit (**Fig. 2C**).

**Figure 2B.**
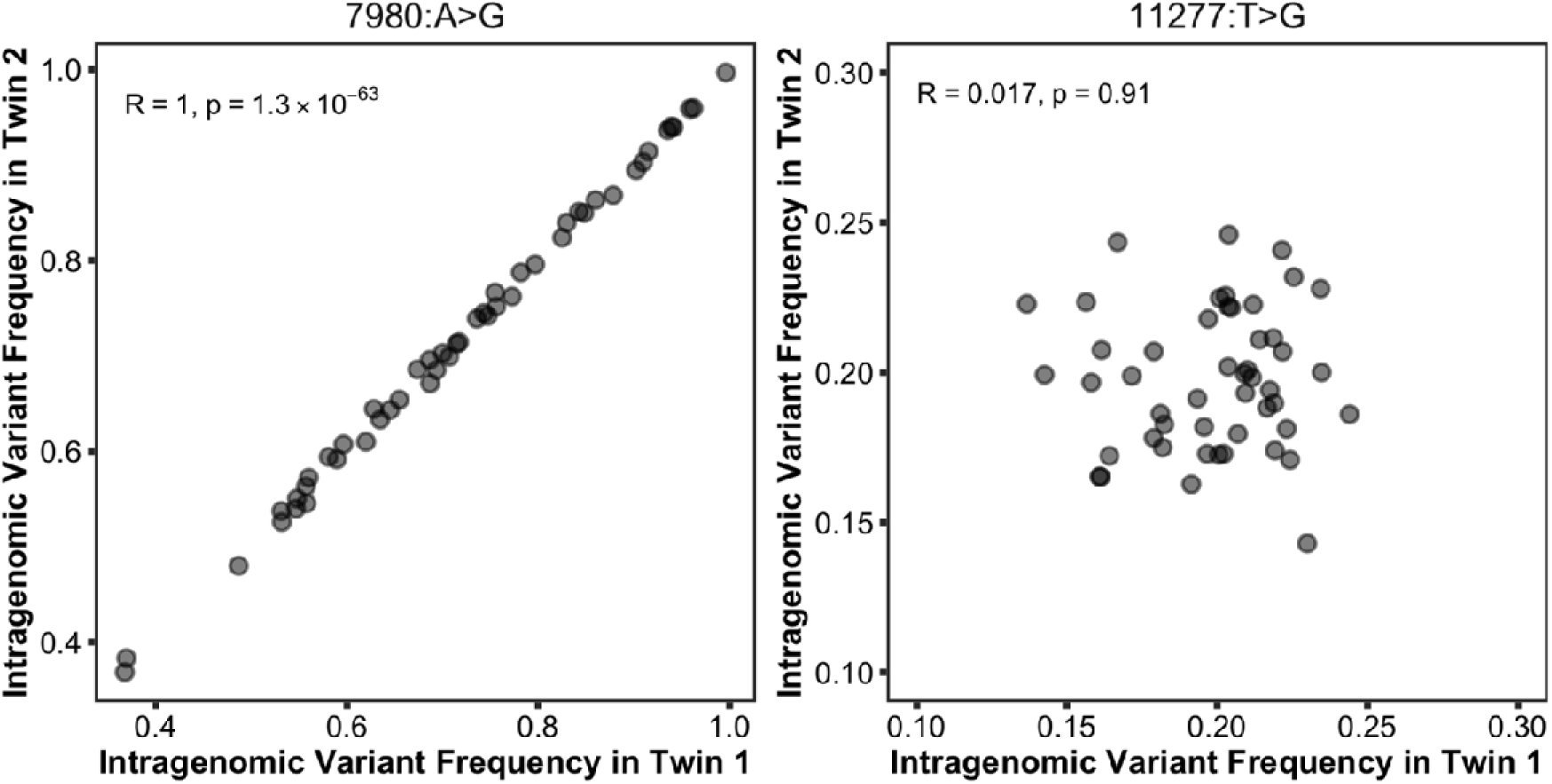
Examples of intragenomic variant frequencies comparisons in MZ twins. The left panel shows an rDNA variant with high intra-twin-pair correlation, whereas the right panel shows a variant with extremely low correlation.

**Figure 2C.**
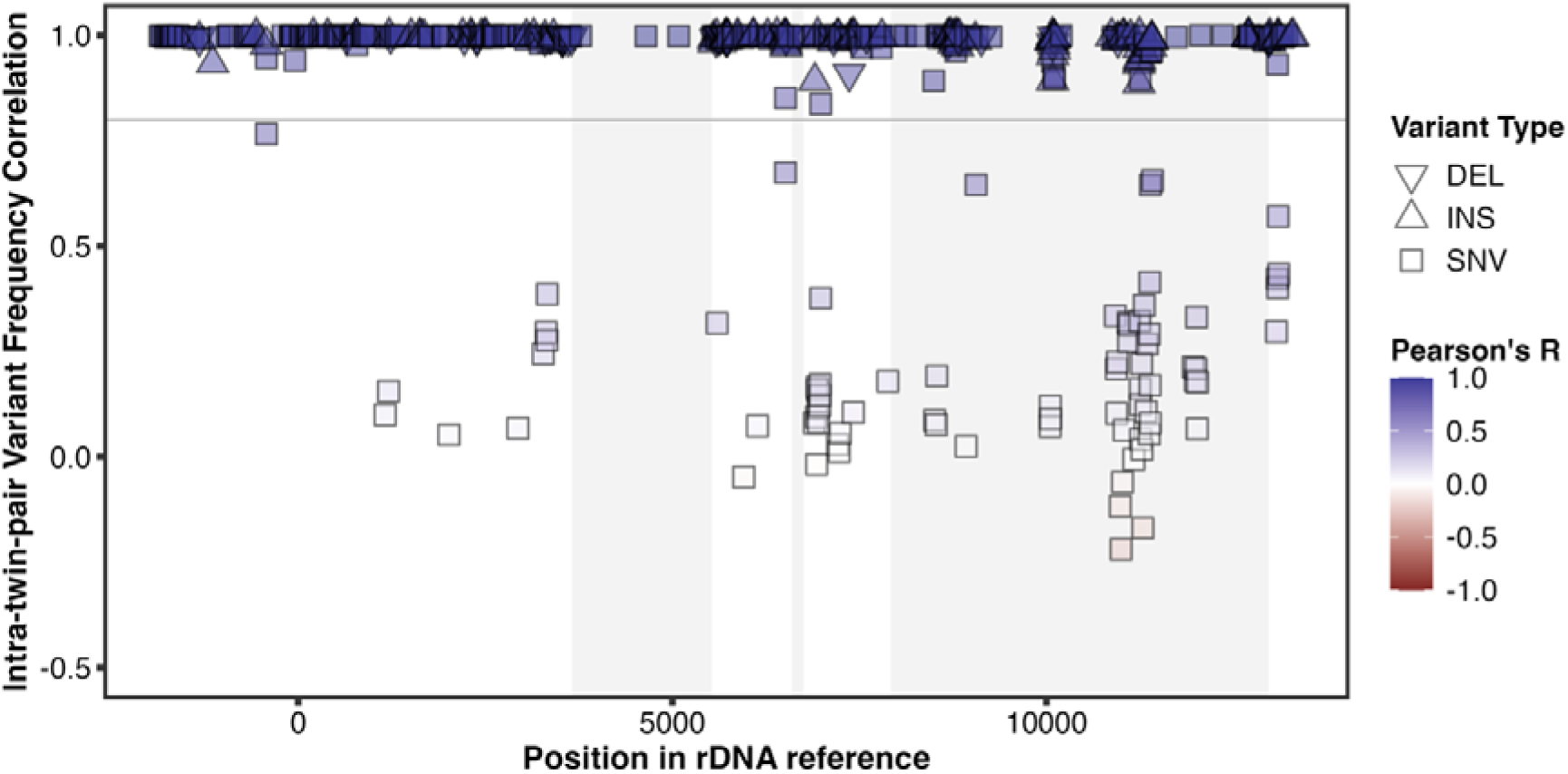
Intra-twin-pair correlation in estimated variant frequencies across the rDNA transcriptional unit. The horizontal line indicates the R = 0.8 threshold chosen.

The apparent lack of intra-twin-pair correlation in some variants could potentially arise due to somatic variation, causing the rDNA units of individuals within each twin pair to diverge throughout their lifespans. This phenomenon could occur due to different allele-specific changes in the total number of rDNA CN, most likely in the form of CN loss^[25]^. Our previous research in the UK Biobank^[6]^ alongside other observations^[26,27]^, however, suggests no such loss of rDNA copies with age occurs in humans. Alternatively, point mutations could accumulate across rDNA units during an individual’s lifespan to the extent that the relative proportions of alleles substantially diverge between MZ twins later in life. The fact that every single variant with R < 0.8 is an SNV could support this interpretation (**Fig. 2C**). Moreover, virtually all SNVs below the R < 0.8 threshold correspond with the same base change (T>G, or, equivalently for the other strand, A>C; **Fig. 2D**). This may suggest some particular context where somatic mutations are more likely.

**Figure 2D.**
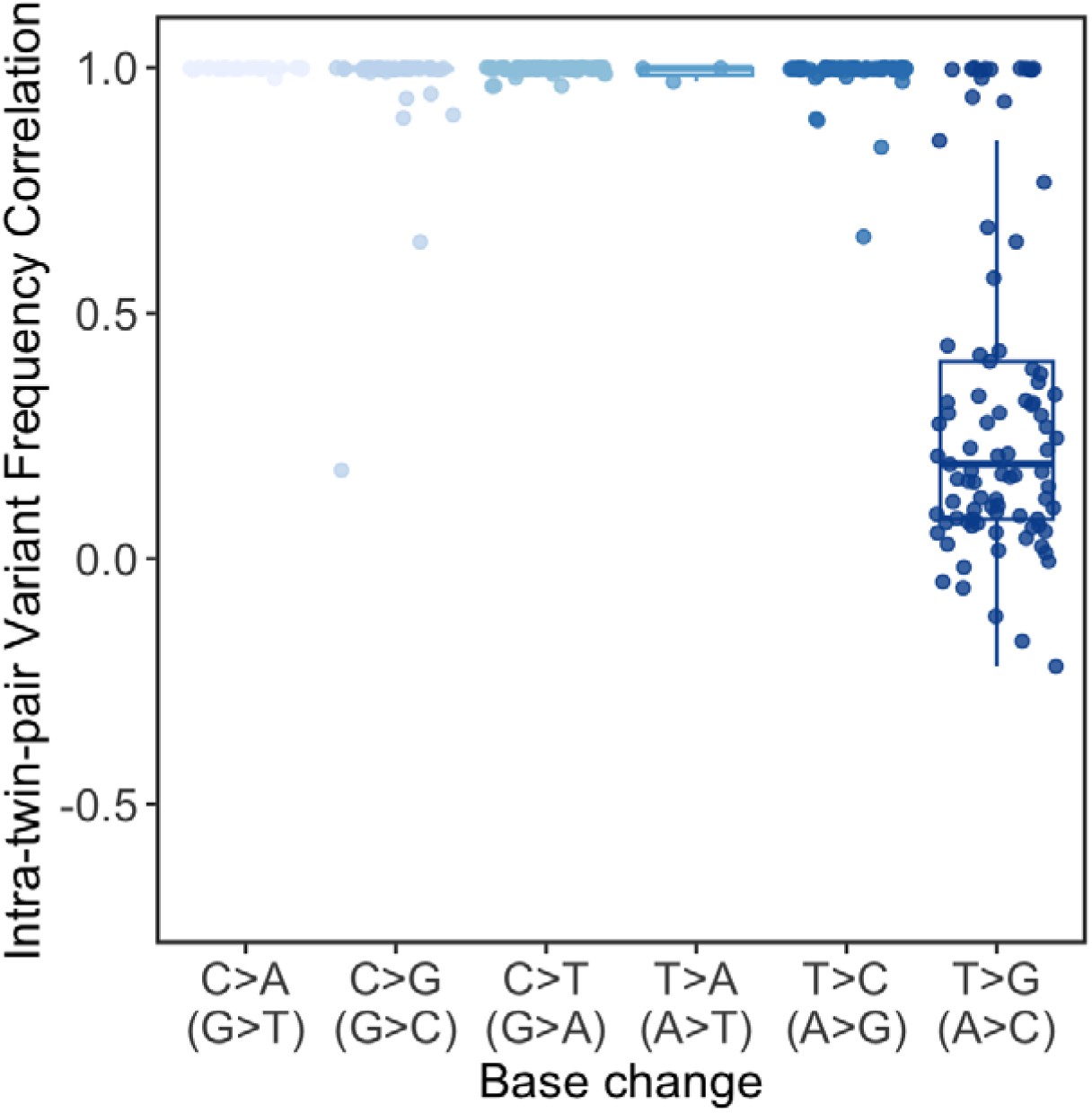
For the SNVs in Fig. 2C, relationship between the intra-twin-pair correlation in estimated variant frequencies and the change in nucleotide corresponding to the specific variant. The changes in parentheses indicate that complementary changes were grouped together.

A more plausible explanation, however, is that the apparent differences between MZ twins are technical artefacts. When comparing short-read Illumina and long-read Oxford Nanopore (ONT) data for the same 1000 Genomes Project GBR individual, it is clear that the intragenomic frequencies for variants with R < 0.8 in UKB MZ twins are virtually undetected in ONT data, but the two technologies agree for variants with R ≥ 0.8 (**Fig. 2E, fig. S2**). This suggests that the Illumina base calls for actual Ts may be reported as Gs (and, equivalently, As reported as Cs) at levels that do not affect conventional low-ploidy variant calls but may cause false positives in highly repetitive parts of the genome where variants are interpreted as an intragenomic frequency continuum. Illumina’s 2-Channel sequencing by synthesis (SBS) technology employed in their current high-throughput devices might help explain this effect, with the high GC content of some specific rDNA regions potentially exacerbating the issue. We thus discarded all variants with intra-twin-pair correlation below 0.8 for any further analysis, thus leaving 380 variants for further consideration.

**Figure 2E.**
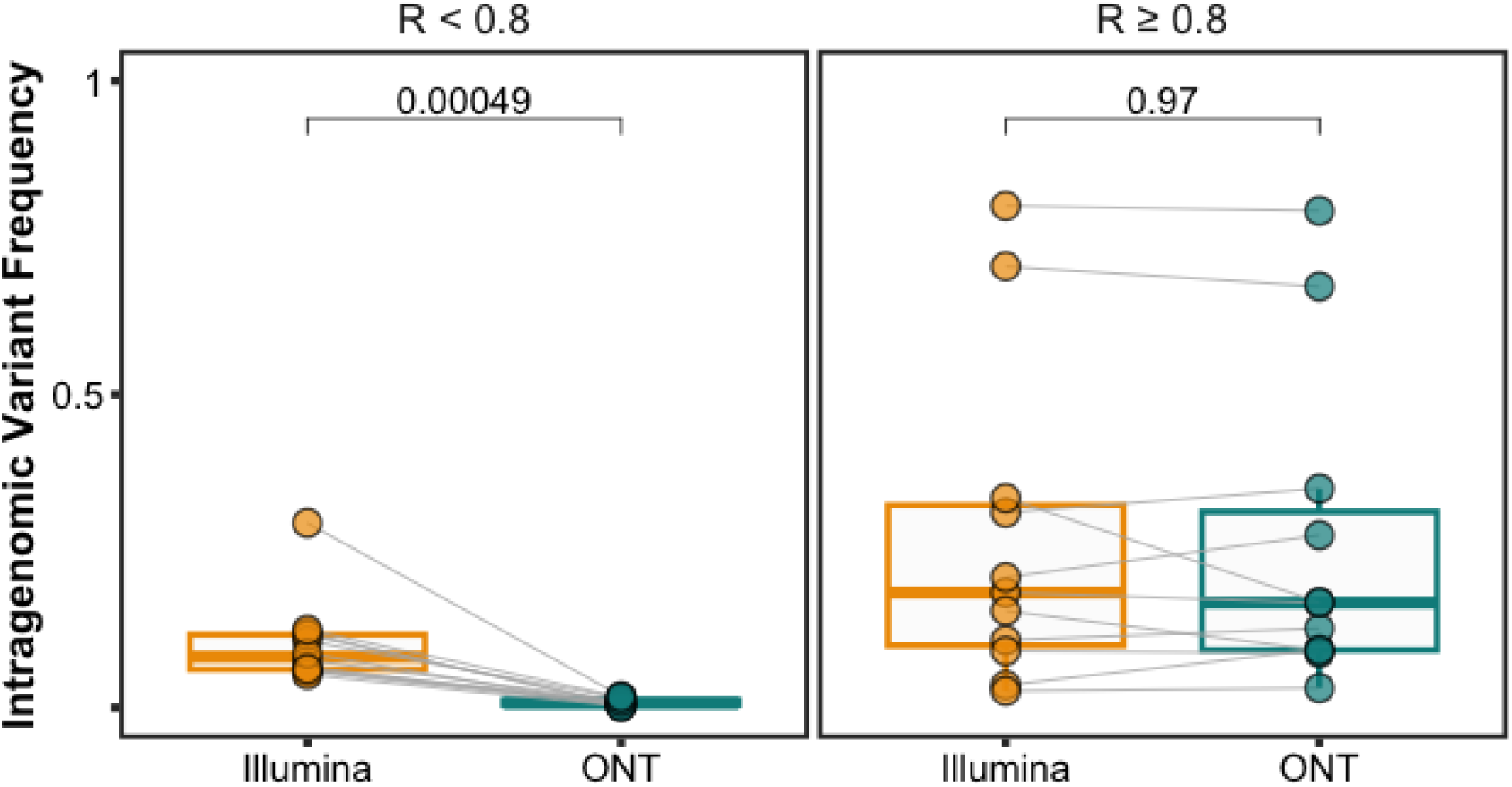
Comparison between the 28S variant frequencies detected from Illumina short-read and Oxford Nanopore (ONT) long-read for HG00127, a GBR participant from the 1000 Genomes Project, split by the intra-twin-pair correlation of each variant obtained on UKB MZ twins.

#### Extracting rDNA analogues from Hg38 has minimal impact on variant frequencies

Since only a relatively small portion of each sequencing library comes from the rDNA, processing solely potential rDNA reads has an enormous impact on the computational efficiency, and thus feasibility, of the analysis. Our previous rDNA CN proxy estimates relied on identifying regions of the Hg38 assembly where 18S reads map, showing that extracting reads from those regions can equate to full realignments of complete sequencing libraries^[6]^. For the current analysis, however, not only is the region of interest larger than the 18S, but also substantially more variable, which could lead to rDNA reads being mapped to further regions of Hg38. Biases in the mapping locations of various alleles of the same variant position might help explain differences in the variants detected in previous studies, with some well-known pervasive intragenomic variants being fully absent due to only a subset of rDNA analogues having been considered^[3]^.

To reduce the risk of missing relevant rDNA analogue regions, we first generated synthetic Illumina reads from the rDNA consensus reference and observed where they mapped when aligned to the Hg38 assembly. We also examined the realignments to the Hg38 plus rDNA tailored assembly reported above to determine where reads now mapping to the rDNA unit had previously been mapped. Both approaches coincide, reporting virtually all synthetic reads and over 98% of the real reads mapping in Hg38 to the same specific windows neighbouring the 18S analogues employed for the CN analysis (**fig. S3**). Furthermore, comparing the variant frequencies obtained from full realignments with those from prior extraction of these putative analogue regions in the MZ twin data confirms the validity of this latter approach. Most variant positions reach identical frequency estimates in either case, with only two variants having Pearson’s R below 0.8 (**Fig. 2F**). In both cases, the decrease in correlation appears to arise due to the variant not being detected at all in a small subset of individuals and not necessarily due to systematic biases in the estimates that could suggest missing analogue regions (**fig. S4**). Regardless of the underlying reason, we only retain variants with R > 0.8 between full realignments and prior analogue extraction.

**Figure 2F.**
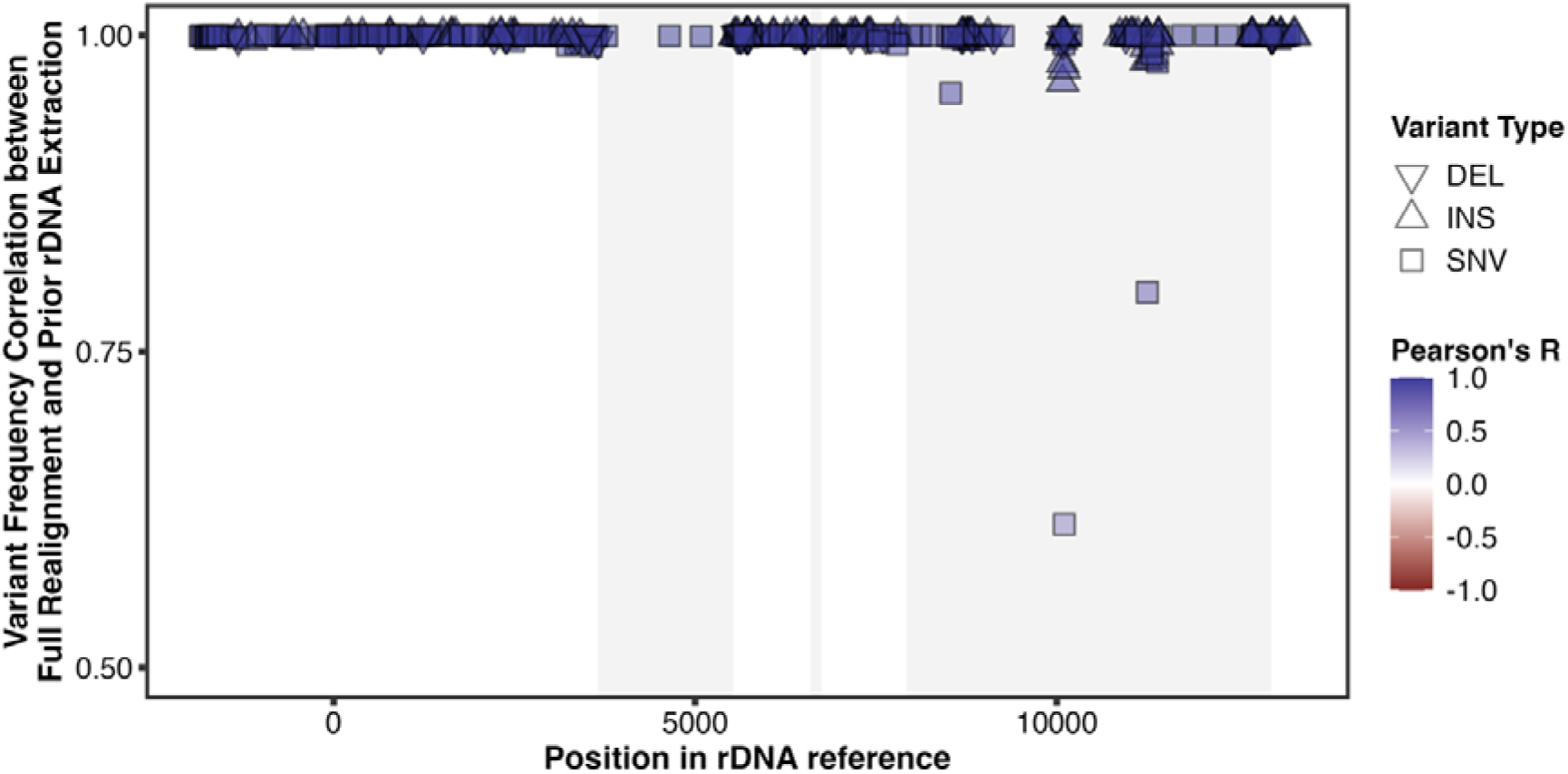
Correlation between variant frequencies across the rDNA transcriptional unit estimated from full library realignments and the prior extraction of rDNA analogue reads.

#### Hundreds of rDNA sequence variants are suitable for UKB phenotype association analyses

The selection criteria detailed above leads to 378 sequence variants we deem suitable for further analysis (**Table S1**). These are located across 317 distinct positions between -1848 (upstream of the TSS) and 13294 (within the 3’ ETS) of the KY962518.1 rDNA consensus reference, and correspond to 217 SNVs and 161 INDELs, 48 of which are reported as deletions and 113 as insertions when compared to said reference. As expected, the spatial distribution of the variants is not uniform, with higher density of variation throughout the promoter and spacer regions compared with the rRNA genes, and with the 28S also harbouring substantially more variation than the 18S or 5.8S (**Fig. 2G**). Regarding the intragenomic frequencies, we can clearly distinguish variants for which virtually all twin pairs harbour at least two alleles from those for which most pairs show no variation whatsoever and only a few some degree of biallelism, with usually the reference allele clearly dominating (**fig. S5**) The well-known variant 60 bp into the 28S sequence^[28]^ (here reported as 7980:A>G) belongs to the first group, whereas the small subset of detected 18S variants belong to the latter, with the vast majority of twin pairs displaying no 18S variation (**fig. S6**).

**Figure 2G.**
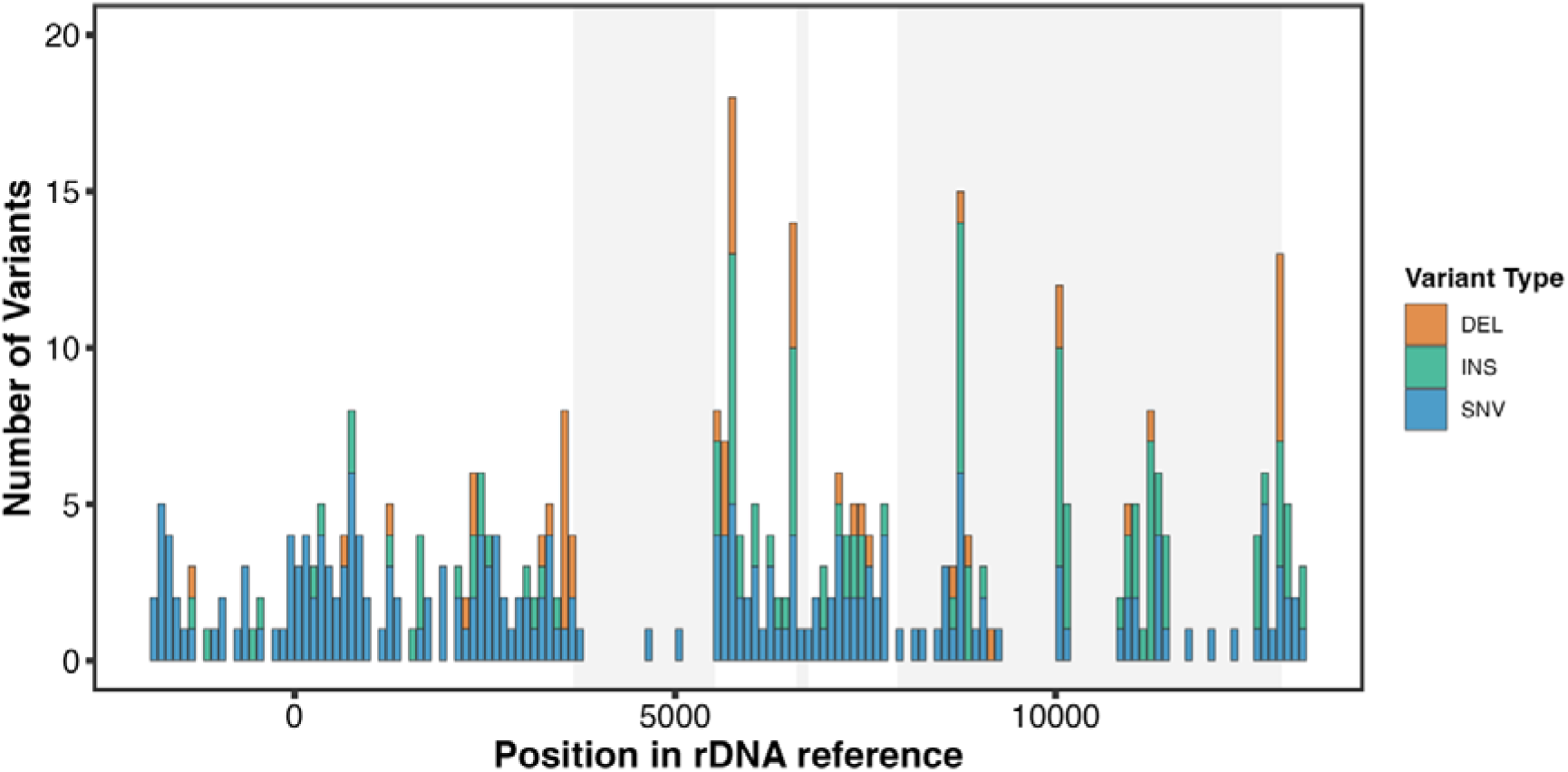
Positional distribution across the human rDNA transcriptional unit of variants selected for further analysis.

### Intragenomic rDNA variant frequencies associate with human traits in UKB

#### 28S intragenomic variant frequencies often associate with UKB phenotypes

For each of the 378 variants selected in the previous section, we then asked if their intragenomic variant frequency associates with phenotypic traits. To this end, we obtained frequency estimates for 490,398 UKB participants using lofreq^[29]^, but only employed a set of 297,010 unrelated WB individuals in association tests. Tests were then conducted independently for each variant using PHESANT^[30]^, following the same procedure we previously reported for rDNA CN^[6]^ to regress phenotypic traits on rDNA variation, but replacing the 18S Ratios with intragenomic variant frequencies (valued between 0 and 1). We also limited the set of considered phenotypes, with 419 distinct traits for which `PHESANT` provided valid output. This reduced set excludes, among others, fields related with cancer (such as diagnoses and histology results) since large, devoted cohorts including both germline and tumour WGS data exist, which should provide much more powerful insights into the potential role of rDNA sequence variation in cancer.

Out of the 158,382 combinations between the 378 variants and 419 phenotypes in the analysis, 34 associations reach statistical significance at an FDR level below 0.01 (**Fig. 3A**, **Table S2**). All these hits correspond to variants within the 28S. In particular, 17 distinct variants out of 102 (16.7%) belonging to the 28S have FDR-significant associations. Since variants within the 28S have the potential to be expressed in the rRNA and thus be incorporated into mature ribosomes, this suggests rDNA sequence variants may impact phenotypic outcomes through the formation of a pool of heterogeneous ribosomes, as it has long been speculated^[14,31]^.

**Figure 3A.**
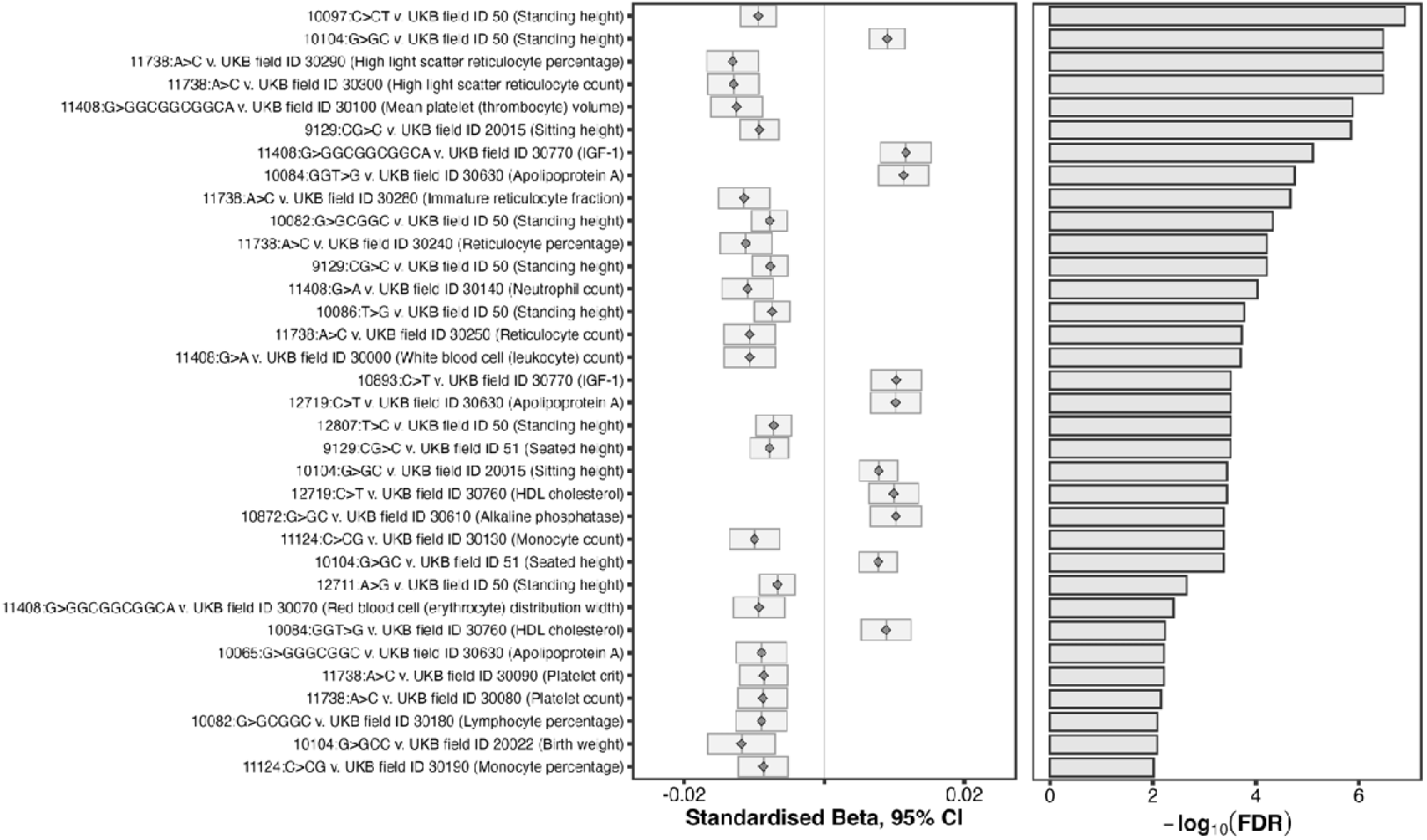
Effect size (Left) and significance level (Right) of associations between intragenomic variant frequencies and UKB phenotypes at global FDR < 0.01, estimated from unrelated WB individuals.

#### A cluster of neighbouring ES15L variants associates with body size-related traits

The top two FDR-significant associations both correspond with the same phenotype ("Standing height") and with variants located less than 10 bp apart. In particular, the intragenomic frequency at 10097:C>CT appears negatively associated with the trait (p = 4.98×10^-13^) whereas 10104:G>GC appears positively associated (p = 4.27×10^-12^). In fact, several variants in the close neighbourhood of these two associate with height and/or other body size- or growth-related phenotypes. This becomes clearer if we depict positionally all associations at variant-level FDR < 0.01 (**Fig. 3B**). A single cluster of variants around position 10100 associates not only with diverse body size measurements (including "Weight", "Waist circumference", and "Birth Weight" aside from different versions of "height") but also "Cholesterol", "Apolipoprotein A", and "HDL Cholesterol" — closely associated with weight — as well as "Other intervertebral disk disorders" — which could relate to height.

**Figure 3B.**
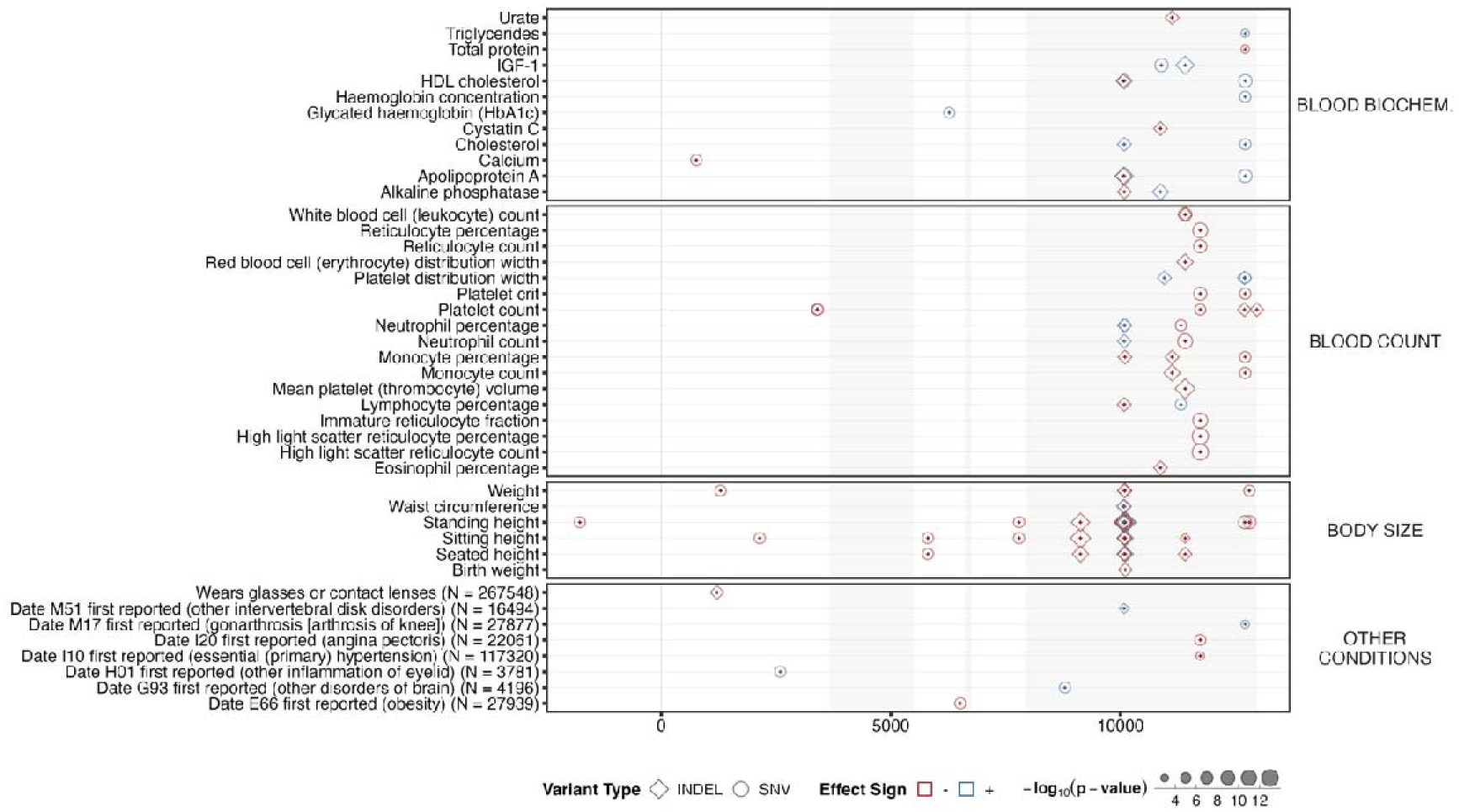
Positional distribution of phenotypic associations across the rDNA transcriptional unit with per-variant FDR < 0.01.

Focusing on the 28S and separating the variants makes it clear that the cluster around 10100 is actually formed by 8 distinct variants at 6 different positions within a span of fewer than 40 bp (**Fig. 3C**). These all fall within the so-called ES15L — the Expansion Segment 15 of the Large ribosomal subunit. In fact, the vast majority of these variants with putative phenotypic associations fall within Expansion Segments (ESs). The exceptions are 12971:CCG>C — which coincides with the 3’ end of the 28S — and 11738:A>C — whose association with multiple cardiovascular- and myeloid-related phenotypes deserves devoted attention.

**Figure 3C.**
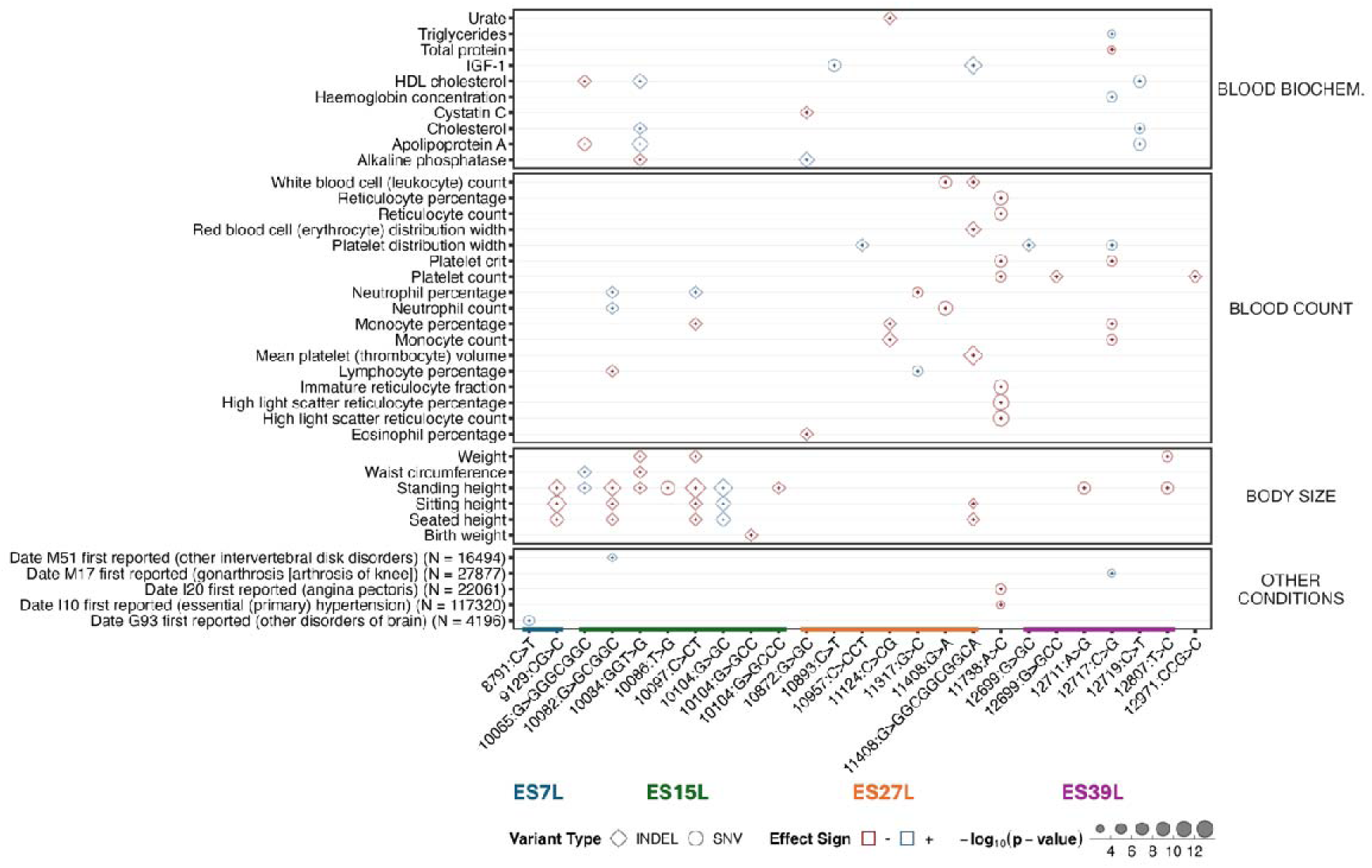
Phenotypic associations across the 28S with per-variant FDR < 0.01. The corresponding expansion segments are indicated.

As noted before in smaller cohorts^[4]^, the correlation between 28S intragenomic variant frequencies in the UKB is substantially higher within ESs, and particularly within ES15L (**Fig. 3D**). ES15L variants were not included in previous attempts to build 28S haplotypes (or "morphs"), but the tight correlation between the variant frequencies within the region suggests they might form local haplotypes. Preliminary analyses suggests that might be the case, particularly when considering the multiple variants at position 10104 separately, with some combination of alleles at positions 10065, 10082, 10084, 10086, and 10097 appearing in a median of around 30% of each individual’s rDNA units (**Fig. 3E**). Multiple other combinations seem to form 15% or fewer of the rDNA units, leading to a landscape substantially less clearly defined than what was previously observed in inbred mice^[23]^. Nevertheless, in the future it might be worth exploring phenotypic associations with the intragenomic prevalence of these variant combinations instead of each one separately.

**Figure 3D.**
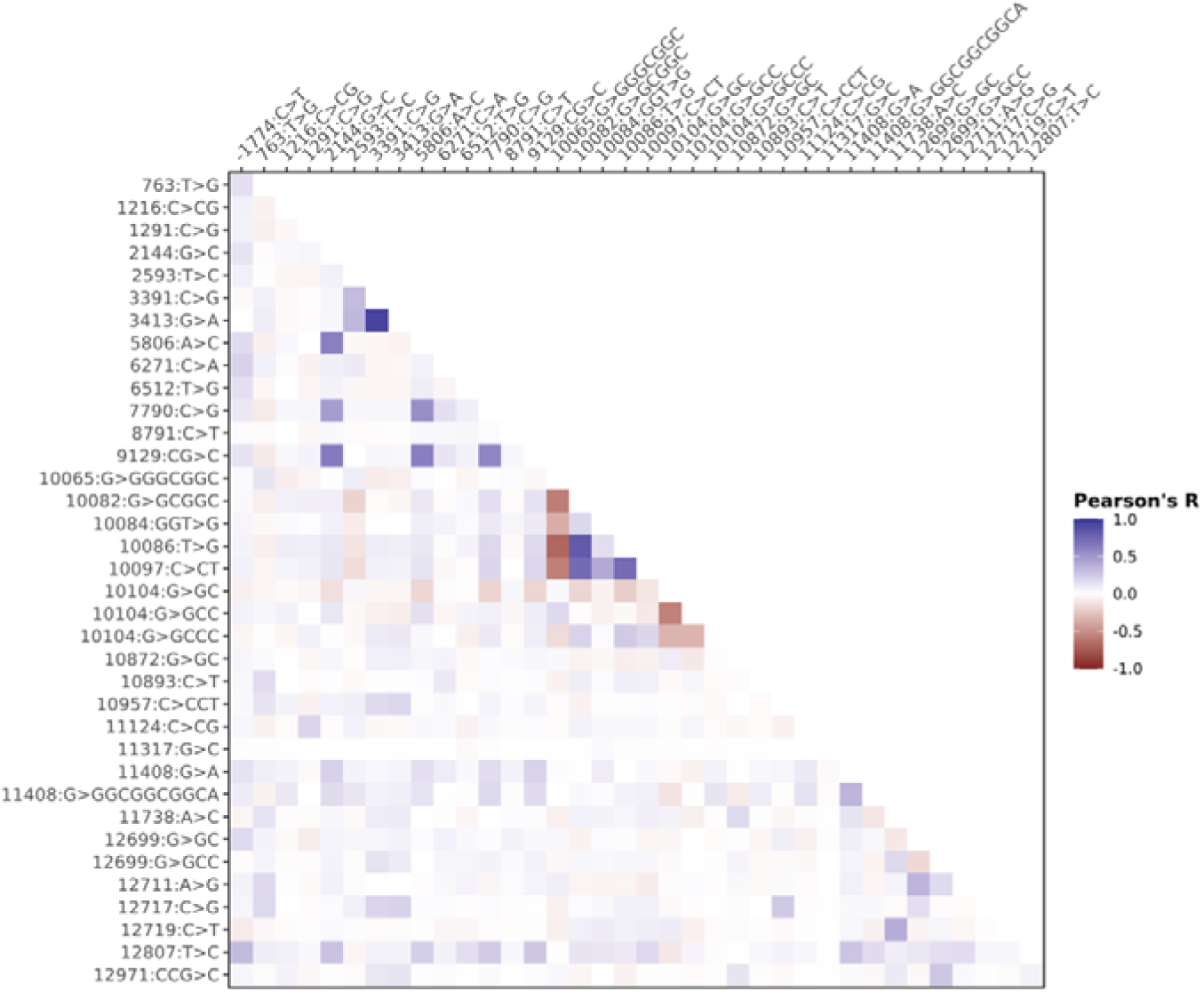
Heatmap of correlations between intragenomic variant frequencies across pairs of variants with variant-level FDR < 0.01 associations.

**Figure 3E.**
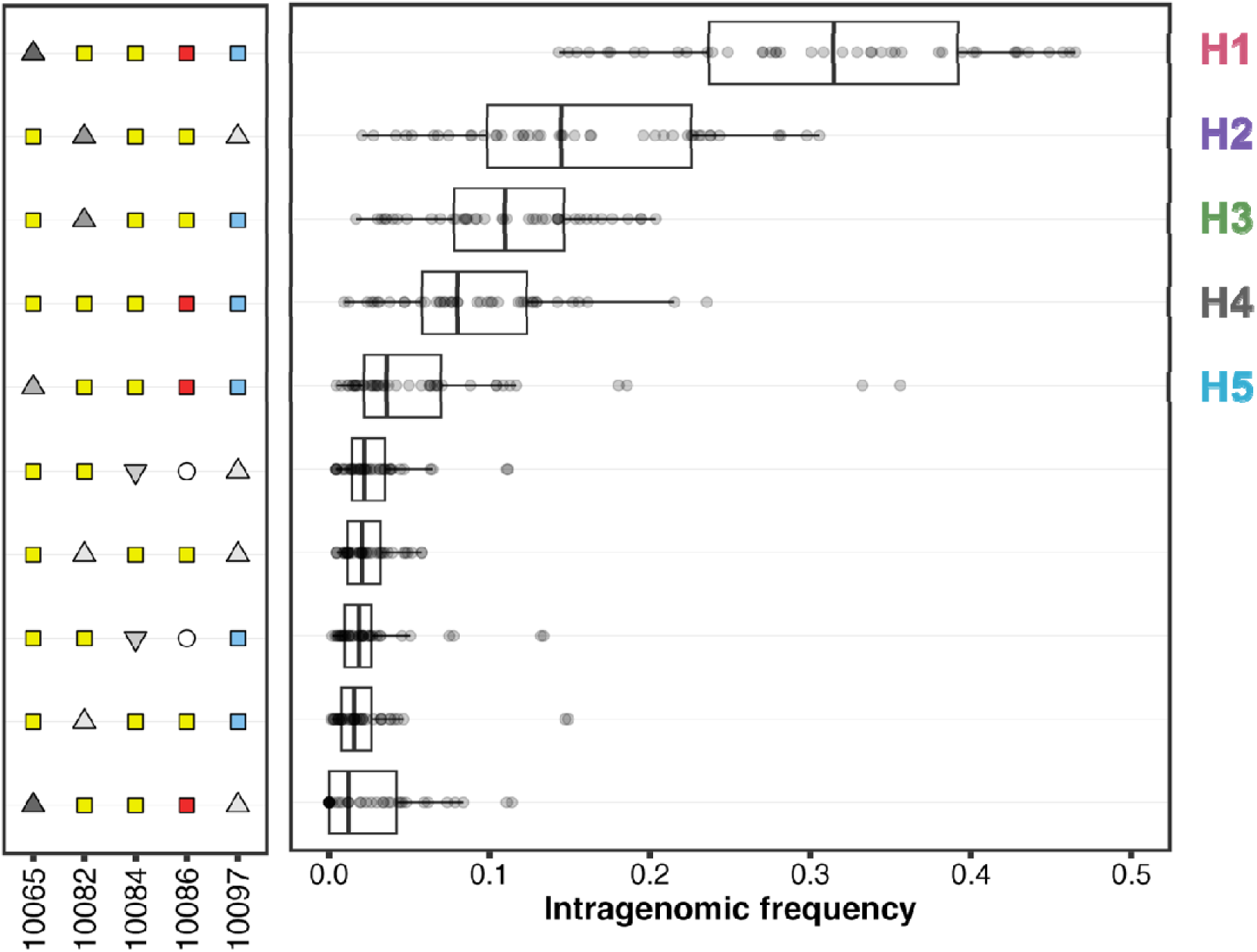
Ten most frequent combinations of alleles at a subset of ES15L variants in MZ twins sequenced by deCODE in the second sequencing release. Squares represent single nucleotides (’C’ is blue, ’G’ is yellow, and ’T’ is red); triangles represent INDELs (upwards being insertions and downwards being deletions, with their darkness representing the size of the INDEL itself); and circles represent absent positions (due to a prior deletion spanning them). The combination labelled as H4 coincides with sequence present in the KY962518.1 rDNA consensus sequence.

#### Phenotypically-associated variants are expressed in GBR individuals from the 1000 Genomes Project

ESs are inserted regions within the eukaryotic rRNAs, greatly increasing their size compared with the prokaryotic sequences. They tend to locate as protrusions in the surface of the mature ribosomes (**fig. S7**). These regions harbour most rRNA variation across and within species but their function is still poorly understood^[4,32]^. It is believed that they may contribute to modulating translation by binding ribosome-associated proteins^[33]^. Moreover, it has been noted that 28S ESs partly resemble mRNA sequences, particularly ES15L, which also appears to have been greatly expanded in mammals, particularly in hominids^[34]^. Similar to what others have previously reported^[4]^, the variants we observe in ES15L tend to affect the number of GGC tandem repeats in the region. Comparisons of 2D and 3D structural models of ES15L suggests that the rRNA conformation changes across observed combinations of variants in the region (**Fig. 3F**, **fig. S8**). This can thus impact how the ES protrudes out of the ribosome’s surface and, in turn, affect protein and/or mRNA binding, potentially influencing phenotypic outcomes through differential modulation of translation.

**Figure 3F.**
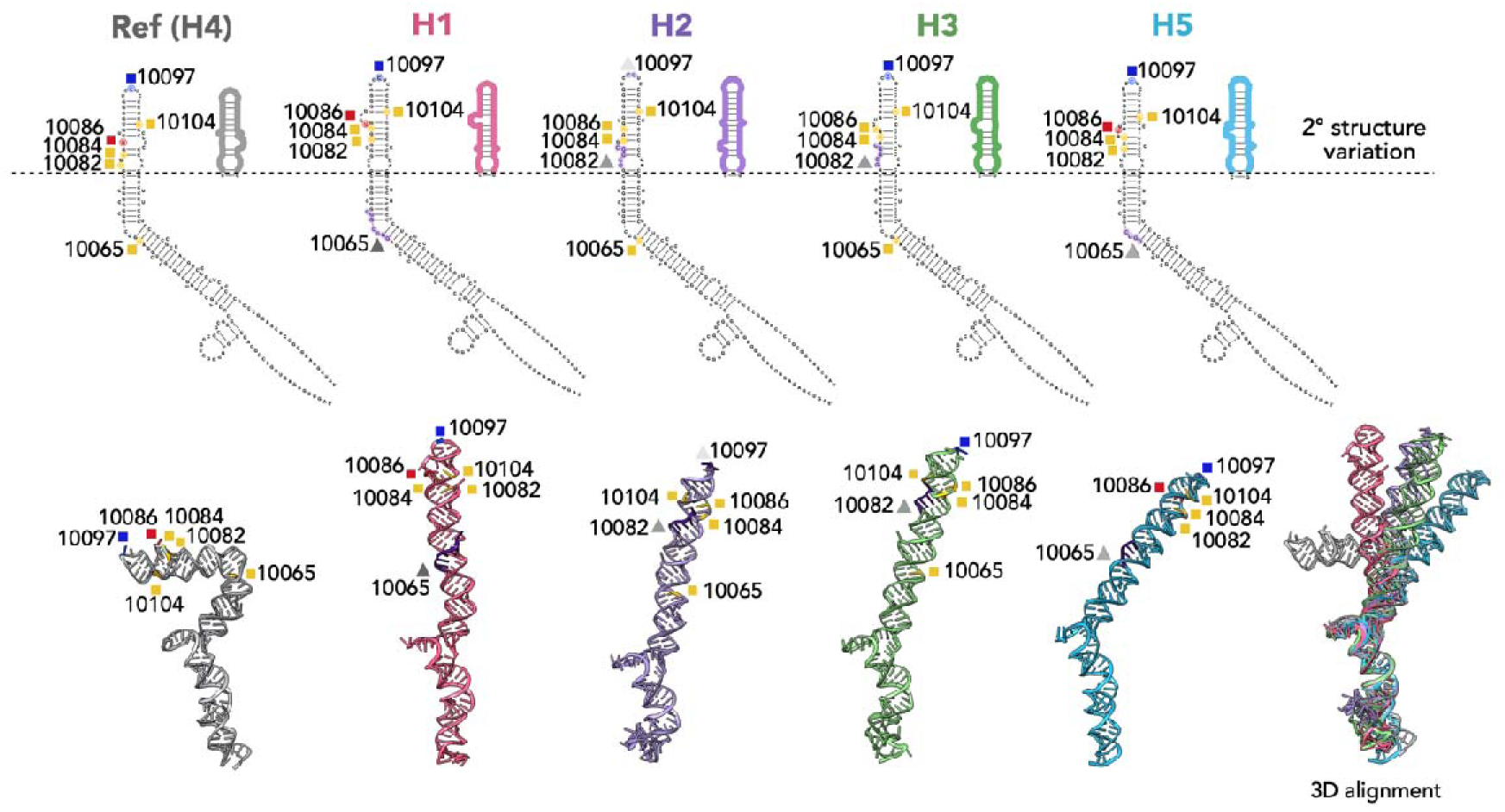
Predicted secondary (Top) and tertiary (Bottom) structure RNA models of the 5 most frequent combinations of alleles at a subset of ES15L variants. Symbols and colours indicate SNVs and INDELs as in Fig. 3E. Changes to secondary structure are observed above the dashed line. Corresponding 3D structures are given beneath the 2D maps. Alignment of all 3D structures to the reference (grey) reveals potential differences in variant conformations.

This hypothetical mechanism, however, becomes irrelevant unless the observed variants are expressed in rRNA. Unfortunately, the UKB does not yet provide RNA-Seq data. Instead, we obtained publicly-available RNA-Seq data for 28 GBR participants and calculated variant frequencies at the positions of interest. We observe substantial expression of most of the ES15L variants with phenotypic association, from a median of 7.3% in 17/28 individuals for 10084:GGT>G to a median of 35% across all 28 participants for 10104:G>GCC (**Fig. 3G**). Nevertheless, similar to what we previously observed in inbred mice^[23]^, the expression levels may not be entirely determined by the rDNA variant frequencies, suggesting allele-specific silencing might exist. Comparing the frequencies obtained in the region for the 25 GBR participants for which both datasets exist, we see that, among the phenotypically-associated variants, 10084:GGT>G also reaches the lowest Pearson’s correlation between DNA and RNA levels, with an R of 0.648, whereas 10097:C>CT reaches the highest, with an R of 0.907 (**fig. S9**). This suggests epigenetic or transcriptomic data in the UKB could help refine the associations, but it seems clear that the expression of these variants and their potential incorporation into mature ribosomes provides a mechanistic path for their effect on phenotypic outcomes.

**Figure 3G.**
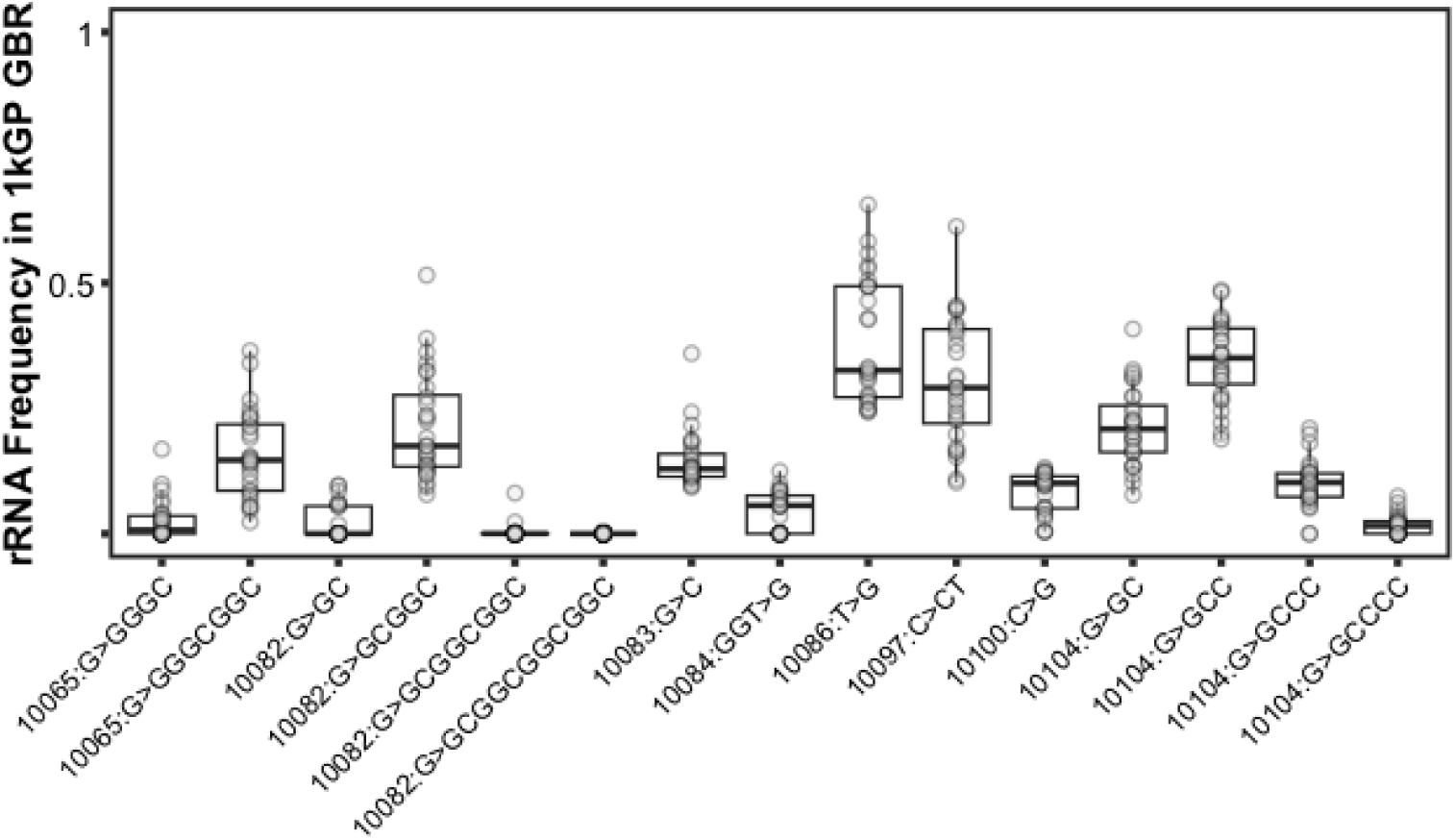
Frequency of expression of diverse ES15L variants in RNA-Seq data from GBR participants of the 1000 Genomes Project.

### Allelic effects are largely independent from total rDNA CN effects

The phenotypic associations identified above for rDNA variant frequencies often overlap the categories we previously reported for total rDNA CN associations, both for blood cell subtype composition and blood biochemistry traits in the UKB^[6]^ and body size elsewhere^[13]^. It is thus reasonable to wonder whether the allelic effects we observe are simply recapitulating the total CN results. To this end, we now consider whether accounting for rDNA CN impacts the allelic effects.

#### Considering both UKB WGS releases jointly reveals further total rDNA CN phenotype associations

The previously published phenotype association results for rDNA CN in the UKB split the participants into a discovery and a validation set according to the sequencing release to which they belonged^[6]^, thus not fully exploiting the breath and power of the cohort. Analysing all unrelated WB individuals (retaining sequencing centre and release as covariates) provides a more exhaustive overview of the phenotypic impacts of rDNA CN (**Fig. 4A**; **Table S3**). In particular, we see that all the previously highlighted associations remain highly significant, including neutrophil count and percentage and blood biomarkers associated with kidney function, such as Cystatin C. It is worth noting that the risk of kidney failure, both acute (N17) and chronic (N18), now both appear as significantly associated at FDR < 0.01 level separately, alongside a number of other conditions.

**Figure 4A.**
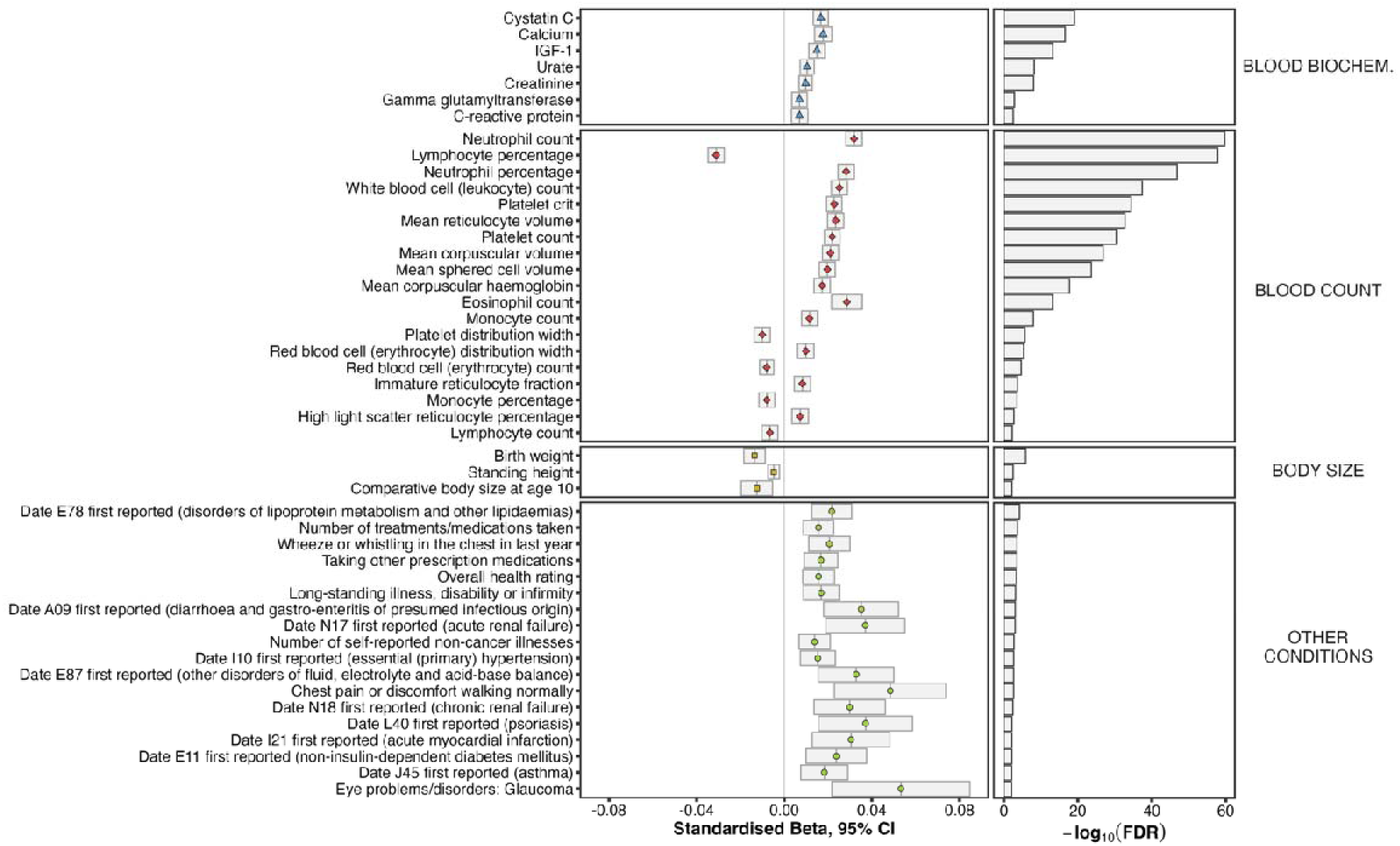
Effect size (Left) and significance level (Right) of phenotypic associations with 18S Ratio (a proxy for total rDNA copy number) in UKB white British participants, combining both sequencing releases. Expanded from [6].

Overall, most associations with total rDNA CN in the UKB are consistent with higher CN being worse for human health, at least later in life. This includes a novel association with "Overall Health Rating" (note — the effect is positive in sign since a lower number represents relatively better "health", "1" meaning "Excellent" and "4" meaning "Poor"). Also novel are the negative associations with three measurements in the "body size" category: "Birth weight", "Standing height", and "Comparative body size at age 10". These further extend the overlap with the allelic associations, but it remains unclear if they involve the same or a different mechanism.

#### Allele-specific rDNA CN effects recapitulate total rDNA CN associations

Having both estimates for the total number of rDNA copies each individual harbours (in the form of 18S Ratios) as well as the intragenomic frequencies of a range of variants in the region enables us to determine the extent to which the precise number of rDNA copies harbouring a specific variant allele associates with phenotype (see **Fig. 1B**). For example, given two individuals (say, X and Y) with different total number of rDNA copies (say, CN_x_ = 400 and CN_y_ = 200) and different intragenomic frequencies at a given variant (say, F_x_ = 0.25 and F_y_ = 0.5), they might experience the same contribution to a particular phenotypic outcome because their cells harbour the exact same number of rDNA copies with a variant allele (in this case, 100). We call this allele-specific CN (AS/CN), which we can calculate for each variant of intertest as the product of the intragenomic frequency and total rDNA CN (or, in this case, the 18S Ratio as proxy). These values can then be assessed for their potential association with phenotypes using the same regression models

Comparing the effect sizes obtained in linear regression models using variant frequency as independent variable (**fig. S10**) with those including allele-specific CN (**fig. S11**) clearly shows an increase of the effect magnitudes across throughout the unit for those phenotypes already shown to associate with total rDNA CN (**Fig. 4A**). The effect of the total number of copies appears to overwhelm that of the particular allele. This can be clearly seen in variants such as 11408:G>A (left panel of **Fig. 4B**), where the association between its intragenomic frequency and neutrophil counts is significantly negative (β = -0.0102 [-0.0139 – -0.00650], p = 5.84×10^-8^) but, when calculating the association with its allele-specific CN, the effect becomes positive (β = 0.0124 [0.00865 – 0.0161], p = 7.30×10^-11^), aligning with the effect of the total number of copies. When the effect of total CN is still opposed to the allelic effect but with smaller intensity, such as the case of the "Immature reticulocyte fraction" in 11738:A>C (right panel of **Fig. 4B**), the effect remains significantly negative in both associations with frequency (β = -0.0111 [-0.0148 – -0.00747], p = 2.48×10^-9^) and allele-specific CN (β = -0.00946 [-0.0132 – -0.00590], p = 3.00×10^-7^), but with a pull in the direction of the total CN association.

**Figure 4B.**
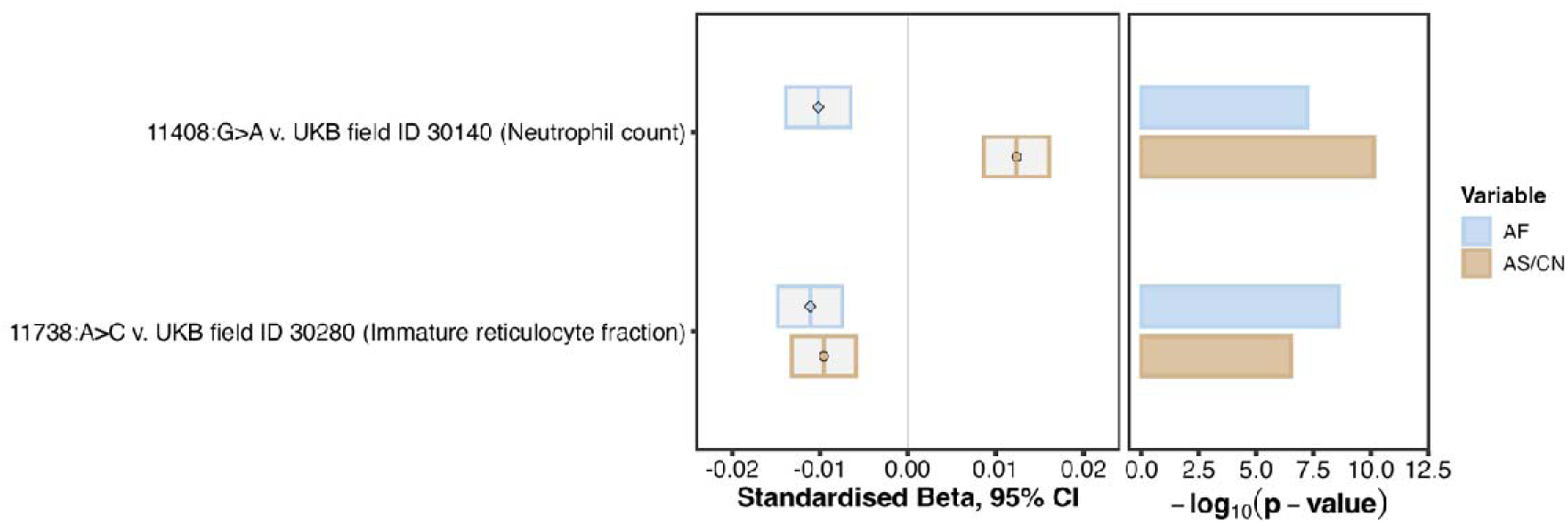
Comparison between effect sizes (Left) and significance levels (Right) obtained for intragenomic variant frequency (AF) and allele-specific copy number (AS/CN) in two combinations of variant and phenotype reaching statistical significance in all cases.

#### Most variant frequency effects remain unaltered when accounting for total rDNA CN

The results above show that allelic and CN effects can oppose each other but, when both components are considered, the associations with CN tend to dominate. It is thus fair to wonder whether incorporating total CN as covariate in the regression models might completely erase any potential association with variant frequencies. If we calculate the difference in the effect sizes obtained both with and without CN as a covariate, we see that this is not the case, with virtually negligible differences in most variant-trait combinations (**fig. S12**). The most salient exception is in 11317:G>C, where including CN as covariate has a marked impact. This is probably due to the substantially larger correlation between the variant frequency at that position and total CN (Pearson’s R ≈ -0.2) than any other considered variant (**fig. S13**). In the particular case of the associations between body size measurements and the frequency of ES15L variants highlighted above, we see virtually no change in both effect sizes and significance levels (**Fig. 4C**). This suggests that the observed associations in the region are independent from the total number of rDNA copies that each individual harbours.

**Figure 4C.**
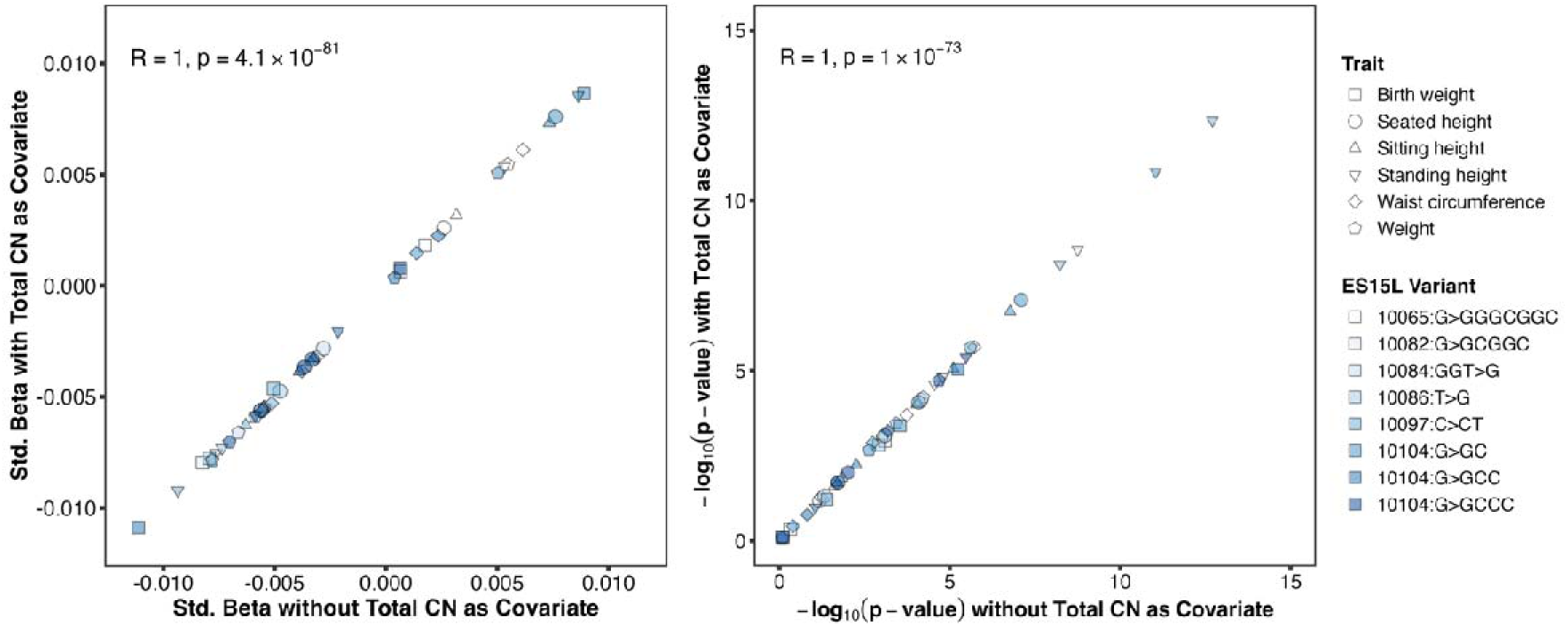
Comparison between effect sizes (Left) and significance levels (right) obtained for all phenotype-associated variants in ES15L with and including total rDNA copy number as covariate in the regression models for body size phenotypes.

## Discussion

Here, using a stringent variant identification strategy, we have shown that germline SNVs and INDELs within the rDNA, a region that has largely been ignored in human association studies to date, are associated with a variety of human complex traits. A key challenge now is to elucidate the mechanistic details of how rDNA genetic variants influence genome function. Inter-individual genetic variation of SNVs and INDELs in the rDNA has been observed in all organisms examined in this regard to date, and in bacteria it has already been demonstrated that rDNA genetic variants influence translational outcomes^[17,18]^. Could similar mechanisms be operating in humans? It is now recognised that the ribosome displays extensive compositional heterogeneity within an organism^[35]^. Potentially, this could dynamically influence translational activity during normal development and disease^[16]^. However, to date, most work on ribosomal heterogeneity has focused on proteins that associate with the ribosome, and the impact of rDNA (rRNA) genetic variation has been much less studied. It could be postulated that incorporation of genetically different rRNAs could impact binding of either proteins or even mRNAs. In the latter case, intriguingly it has been observed that expansion segments extensively match the 5’ ends of mRNAs^[34]^, although it remains to be determined if and how this influences ribosomal function. Regardless, our results provide an important starting point for further investigation into the role of the enigmatic expansion segments. Of course, the rDNA genetic variation might impact other aspects of ribogenesis or ribosome function including rRNA modifications^[36,37]^.

To address the mechanistic details, and limitations of our study, there are several lines of research that can be followed in the future. First, our stringent selection approach from just 49 twin pairs means that we have only considered common rDNA variants. WGS data from larger collections of monozygotic twins or much larger datasets that provide both short and long-read sequencing data will allow robust identification of additional rDNA variants. Second, exploring additional biobank-level datasets will be required to ascertain the impact of rDNA variation in other ancestries and diseases. Third, integrating epigenetic information and rRNA profiling will provide important insights into the impact of rDNA genetic variation on molecular outcomes, especially if profiled in a range of tissues or developmental stages. This should also be combined with methods such as Ribo- or Polysome-Seq to measure the impact on translational efficiency. Fourth, single-cell and/or single molecule analyses will allow more direct associations between rDNA variation and ribosomal output, and provide a measure of somatic heterogeneity of rDNA variation, should it exist. Finally, the most powerful approach would be the ability to genetically modify rDNA genetic variation and combine this with a range of molecular and structural studies. Currently, however, such a genetic editing approach is to the best of our knowledge not feasible for the rDNA.

Understanding the relationship between genotype and phenotype is one of the principal challenges in biology. Despite the highly conserved nature of rDNA, we have shown that germline genetic variation in the region has phenotypic consequences. This variation is not captured by previous studies and represents an important new avenue to fully explain how genotype influences phenotype.

## Methods

### General statistical analysis

Unless otherwise specified, all statistical analyses were conducted using in-house `R` scripts (version `4.3.1`) relying on the data manipulation and visualisation packages from the `tidyverse` version `2.0.0`^[38]^. Differences of means were tested using Wilcoxon signed-rank tests from `R`’s `wilcox.test()` function, including the `paired = TRUE` option when necessary. Correlation coefficients and corresponding p-values were derived using Pearson’s method from the `cor.test()` function.

### Variables of interest and covariates

Phenotypic data and sequencing metadata for UKB participants included in the association models was obtained as previously reported^[6]^. In particular, Sex was obtained from UKB field ID 31, with “0” representing females and “1” males. Age at recruitment was obtained from field ID 21003, with Age Squared derived as the value multiplied by itself. Sequencing Centre information was obtained from field ID 32051. Assessment Centre recorded in field ID 54, Adjusted Telomere Length in field ID 22191, and the 20 first Genetic Principal Components in field ID 22009, were also included as covariates in all analyses. Participants with a value of “1001” in field ID 21000 and “1” in field ID were identified as "White British".

### Identification of relatives and unrelated UKB participants

Using the genetic relatedness information that the UKB provides for 107,076 pairs of participants, with 147,612 individual participants represented, we identify monozygotic twin pairs and construct a set of "fully unrelated" participants. In particular, entries in the relatedness table with kinship > 0.4 were considered monozygotic twins. From these, comparison between rDNA variant frequencies in relatives were limited to pairs where both individuals were identified as “White British” and their sequencing data was generated by the same centre within the same release. The entries of the relatedness table were also employed to create a set of 297,010 unrelated individuals by keeping one participant from each identified "family", as previously described^[6]^.

### Access to sequencing data and computations in the UKB RAP

Variant frequency estimates were calculated for all WGS samples available for both first and second UKB releases, similarly to how proxy rDNA CN estimates had been calculated previously^[6]^. Alignment files in `cram` format were accessed through the UK Biobank Research Analysis Platform (UKB RAP). Since the second sequencing release was made available, these are stored in subfolders the `/Bulk/GATK and GraphTyper WGS/Whole genome GATK CRAM files and indices [500k release]/` path. To enable parallelisation of the computation, the paths to the `cram` files were split into batches and saved in separate files with up to 10,000 participants each. Reads identified as mapping to rDNA-analogue regions of the Hg38 assembly (see below) were extracted using `samtools`^[39]^. This was launched using a pair of scripts (one local and another remote) via the UKB RAP application called `swiss-army-knife` (SAK) version `4.5.0`. The two scripts communicated through the `dxpy` python library, installed locally on a virtual environment (via `pip install dxpy)`. Multiple SAK instances can be executed including in the submission script the options `-iin="${project}${script_path}"` and `-icmd="bash ’${script}’ ’$i’`, as previously described^[6]^. The remote script then reads calls `samtools` to extract the reads in the region of interest as a `bam` file for each `cram` file assigned to its instance (see "Identification of rDNA-analogue regions" below). The paths to each alignment file were prepended with `/mnt/project/`, which enables their processing without needing to previously copy them. The resulting filtered alignment files were then further processed to obtain the variant frequency estimates as detailed below.

### Location of rDNA-analogue regions

Two complementary procedures were employed to identify the regions of the Hg38 assembly used in the UKB alignments where reads originating from the rDNA clusters map. First, we generated 1,000,000 simulated Illumina paired-end reads of 150 bp each from the KY962518.1 rDNA reference using `mason2` version 2.0.9^[40]^. These were then aligned to the unmodified Hg38 assembly present in the UKB (file `GRCh38_full_analysis_set_plus_decoy_hla.fa`, corresponding to the assembly previously employed in the 1000 Genomes Project) using `bowtie2` version `2.4.5`^[41]^. Read coverage across all Hg38 contigs was then calculated using `samtools depth` from `samtools` version `1.19.0`, whose output was then employed to determine which regions accumulate more alignments. Second, we converted the UKB alignments to Hg38 for the same-centre MZ twin pair samples included in the second WGS release to `bed` format using the `bamtobed` tool from `bedtools` version `2.28.0`^[42]^. We then realigned those same samples exclusively to the KY962518.1 rDNA reference using `bowtie2`. These new alignments were also converted to `bed` format. Comparing the mapping locations on a per-read basis from the two generated `bed` files thus provided the regions where rDNA-mapping reads accumulate in Hg38. The two procedures provided a consistent set of coordinates we then employ to extract rDNA-analogue reads from the UKB alignments (**Table S4**).

### Processing of short-read sequencing data

Retrieved short-read `bam` alignment files were converted back to `fastq` format using `samtools`, and then processed using the `repair.sh` script from `BBMap` version `38.95`^[43]^ including the `-Xmx20g` options to remove singleton reads. The output was then processed using `trimgalore` version `0.6.5`^[44]^ in `--paired` mode and aligned to the appropriate reference for each analysis using `bowtie2.`

Intragenomic variant frequencies throughout the rDNA transcriptional unit were obtained from the `bowtie2` alignments using lofreq version `2.1.5`^[29]^. Similar to the procedure described in our prior work^[23]^, rDNA alignments were quality-annotated using `lofreq alnqual` and then processed with `lofreq call` `--call-indels` `--use-orphan`. The resulting calls were then fed to `lofreq filter -v 2000 -a 0.1` to obtain a set of "high confidence" variants. Additionally, `lofreq call` was also executed including the `-a 1 -b 1 -B --no-default-filter` options, which provides an "exhaustive" list of all observed variant frequencies in a sample. Variants present in the "high confidence" output of at least one MZ twin pair were then retrieved from the "exhaustive" output for each sample. Frequencies of 0 were then imputed for all "high confidence" variants absent from the exhaustive mode.

### 1000 Genomes Project sequencing data

#### Short read sequencing data

We retrieved paired-end 1000 Genomes Project (1kGP) WGS `fastq` files from the `sequence_read` folders within the `gridftp/1000g/ftp/phase3/data/` path for the GBR individuals available through the EMBL-EBI public Globus endpoint. Ignoring those generated using ABI SOLiD technologies and those with insufficient read depth, this represents 84 different GBR participants. The downloaded files were then merged according to their sample and strand of origin.

RNA-Seq data for 28 different 1kGP GBR participants was obtained from the Sequence Read Archive (accession PRJNA851328)^[45]^. Of these, 25 were also present in the retrieved WGS data and were used in further analyses.

Aside from the UKB WGS data, the short-read processing procedure detailed above was employed for both WGS and RNA-Seq data from the 1kGP. Alignments generated from a RNA-specific aligner, such as `STAR`, cannot be processed with `lofreq`, and alternative software tools require specific phased transcripts to quantify. The combination of `bowtie2` and `lofreq`, on the other hand, provides the intended output even if the source of the data is not DNA since no splicing variants are expected in the 47S rDNA loci.

#### Oxford Nanopore sequencing data

ONT data for participant HG00127 from the 1kGP was retrieved on the 12th of March, 2024 in `fast5` format from the repository at https://s3.amazonaws.com/1000g-ont/index.html^[46]^. At the time of writing, however, only files in `pod5` format are present in the repository, having been converted to the most recent ONT file format. For backwards compatibility, the current file format can be reverted to `fast5` format using the `pod5 convert to_fast5` tool provided by ONT^[47]^. The `fastq` files included in the `fast5` were then aligned to the Hg38+rDNA reference using `minimap2` version `2.26`^[48,49]^. Reads mapping to the 28S rDNA were identified from these alignments using `samtools`.

Similar to the procedure reported before^[23]^, variant frequencies in ONT data were estimated from the output table generated by `megalodon` version `2.2.9`^[50]^ `with the` `--outputs per_read_variants` option. Variants within the 28S identified in the UKB MZ twins and detected in the WGS data for HG00127 were provided alongside the list of 28S-mapping reads generated above. Only per-read variant calls reported with probability exceeding 0.9 were retained.

### Phenotypic association models

Phenotypic association screens were conducted using `PHESANT`^[30]^ on the set of 297,010 unrelated WB UKB participants for both rDNA sequence and total CN variation (in the form of the 18S Ratios previously calculated^[6]^). The analysis was restricted to a more stringent set of variables than previously reported^[6]^ (**Table S5**). The selected fields were then processed as `PHESANT` requires, prepending an “x” and replacing dashes with underscores in their names.

The `phenomeScan.r` script from `PHESANT` was executed on `R` version `4.2.2` for each variable of interest. Sex, age, age squared, sequencing centre, assessment centre, the first 10 genetic principal components, and adjusted telomere length were used as covariates, and the `--genetic="FALSE"` option was included, since the principal components were explicitly provided in the covariate table. The script was run for each analysis in 100 parts, which were then combined using the `mainCombineResults.r` script. Overall, `PHESANT` generated output for 429 distinct phenotypes for the CN associations and 419 across the variant frequency associations, although the number varies depending on the particular variant.

Linear regression models using `R`’s `lm()` function were fit to compare the effect size estimates in figures **S10** and **S11**, and the differences in fig. S12. The same covariates described above were included in these models, plus the 18S Ratios as additional covariate when appropriate.

### Construction of ES15L haplotypes

An in-house `python` script called `var_extract` was developed to retrieve the alleles present at a given set of positions for each read mapping in the region. This script was constructed similarly to the previously released allele-specific methylation tool `blink`^[23]^, but extending its functionality to enable the identification of INDELs aside from SNVs. To this end, `var_extract` relies on the `pysam` library to traverse each mapped position in a read and parse the base (or sequence of bases) present at the specific reference positions of interest. The output table can then be processed to determine colocalisation between variants in single reads and thus derive the most prevalent "haplotypes" in the region. The similarity between the per-variant frequencies derived from these per-read extractions and those `lofreq` estimates supports the validity of the extraction procedure (**fig. S14**).

### rRNA structural models

To visualise the potential impact of ES15L variants on its structure, 2D RNA maps were first generated with the RNA 2D Templates (`R2DT`)^[51]^ tool and edited using `RNAcanvas`^[52]^. 3D structures of ES15L variants were predicted with the `trRosettaRNA`^[53]^ deep-learning algorithm, using the RNA sequence and dot-bracket secondary structure produced by R2DT as inputs. Predicted structure PDB files downloaded and visualised using UCSF `ChimeraX` version `1.8`^[54]^. 3D ribosome structure obtained from PDB accession 8QOI^[55]^.

## Supplementary Figures

**Figure S1.**
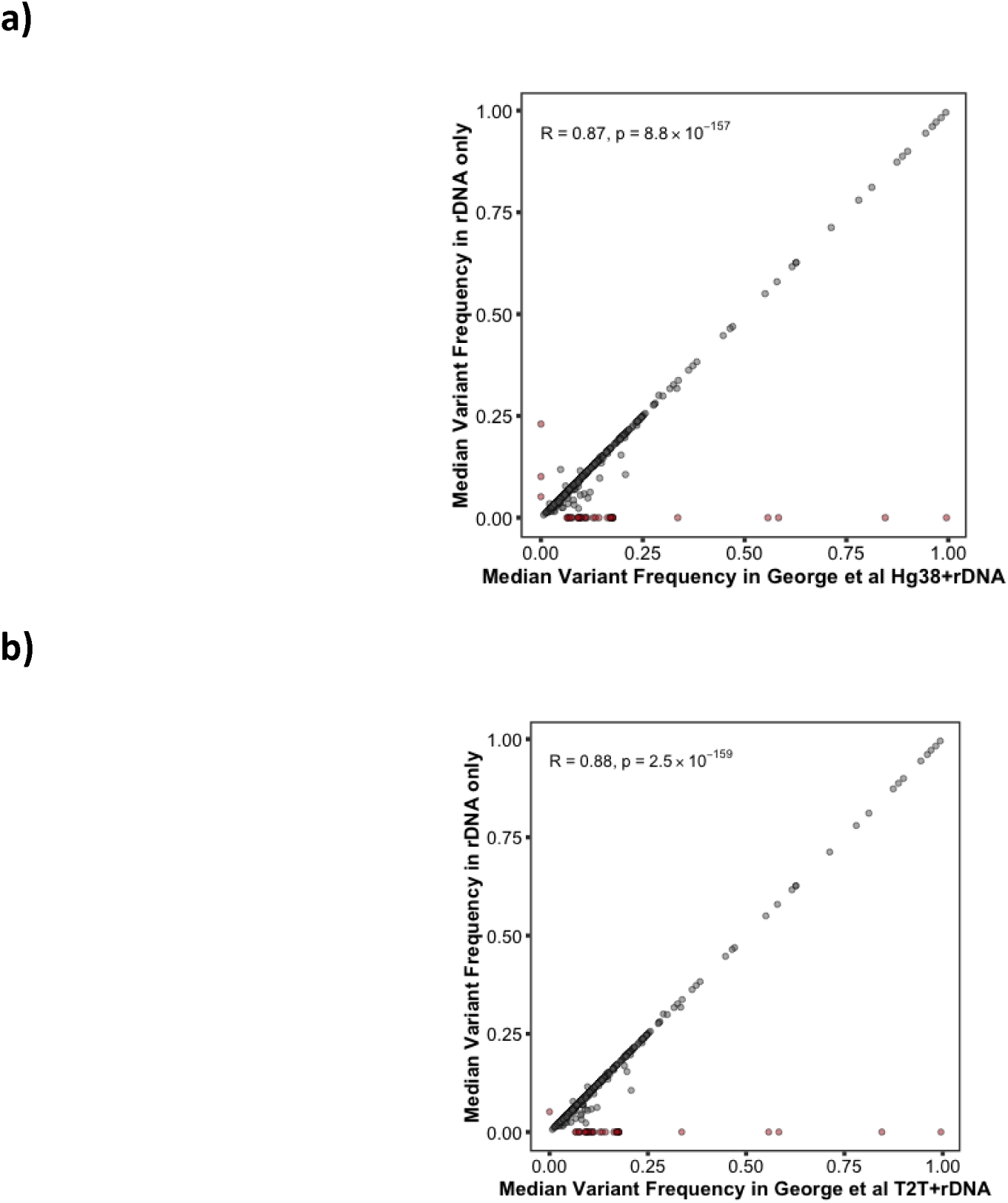
Comparison between the median frequency for each identified variant in the rDNA transcriptional unit estimated from realignments of MZ twin samples from the second UKB sequencing release to full WG+rDNA consensus assemblies generated by George et al^[24]^, and a looped KY962518.1 rDNA reference. Red dots indicate variants that were not detected at any frequency in only one of the approaches.

**Figure S2.**
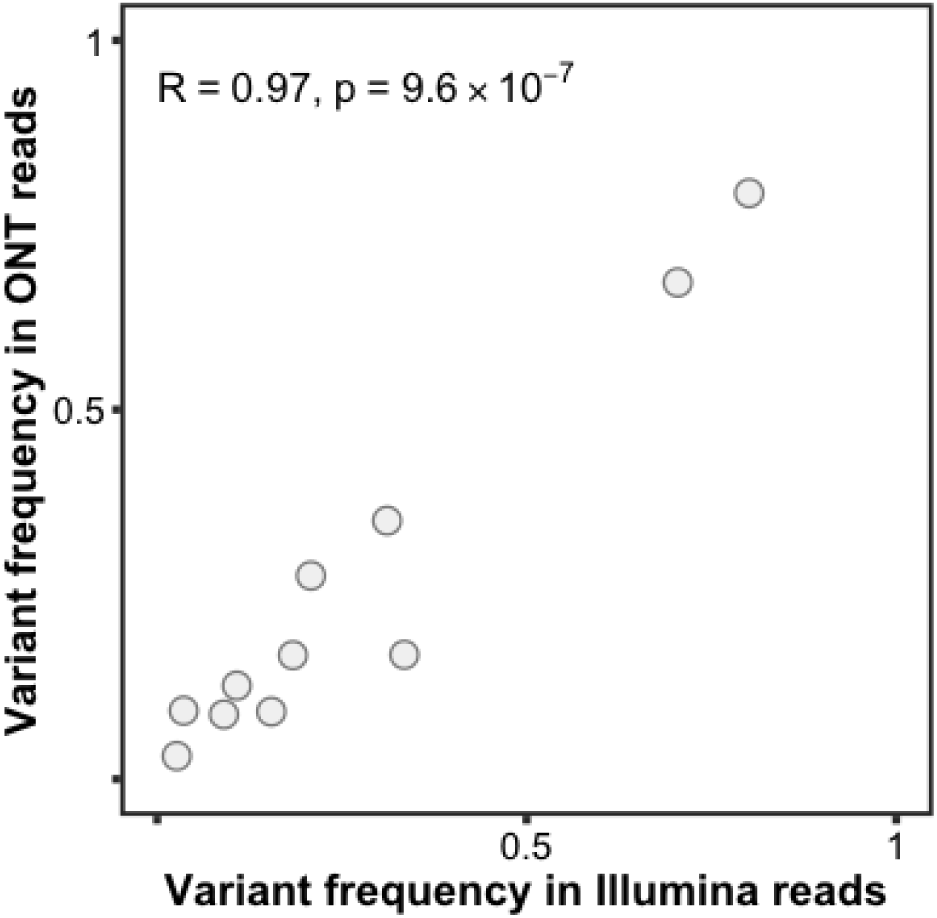
For 28S variants with UKB intra-twin-pair correlation greater or equal than 0.8, correlation between the intragenomic variant frequencies estimated for HG00127 in Illumina short-read and ONT long-read data.

**Figure S3.**
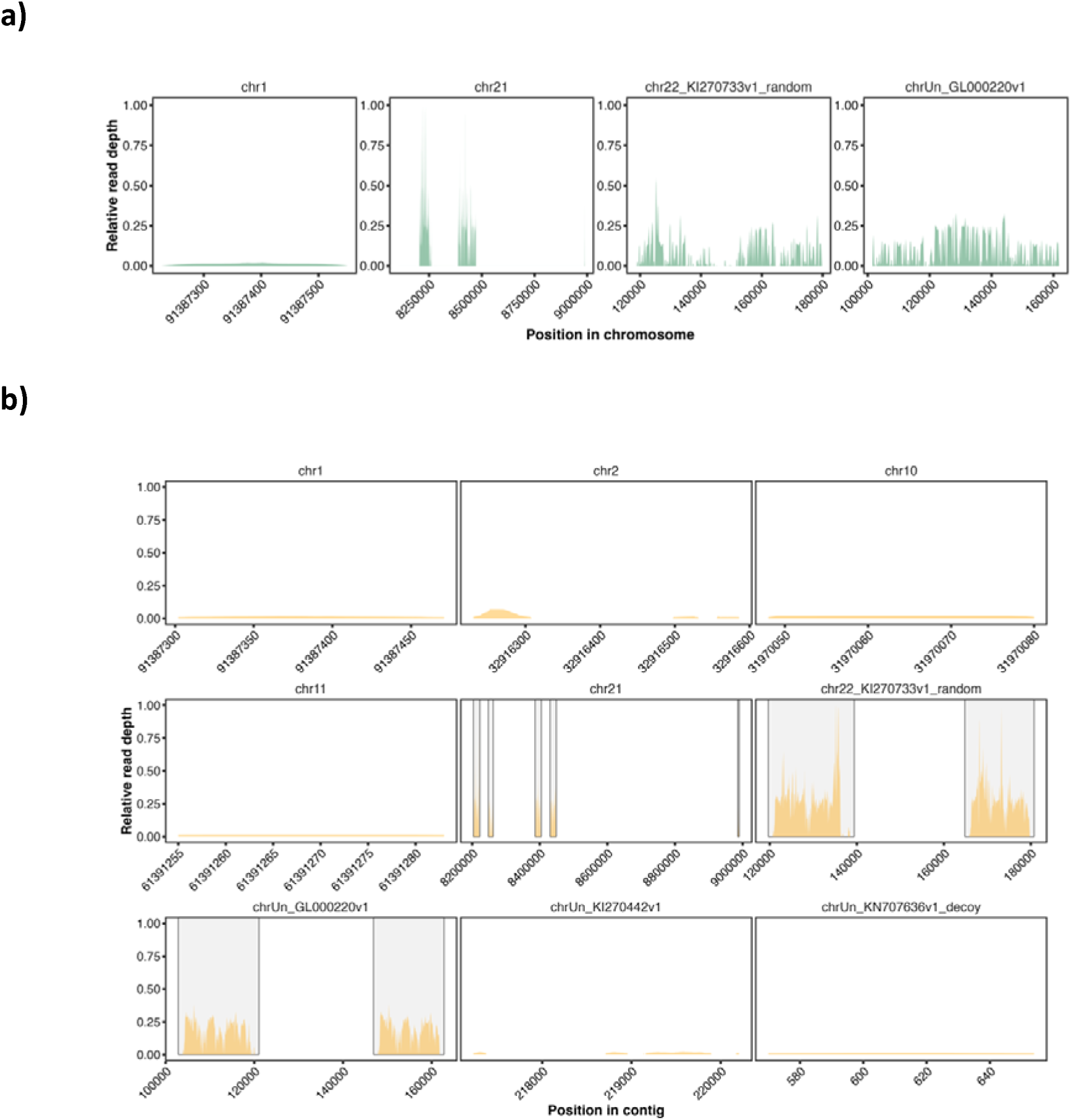
(a) For Illumina reads simulated from the entire KY962518.1 rDNA reference, mapping prevalence across Hg38 contigs where most reads end being allocated on Hg38 alignments. **(b)** For reads that after realignment to a Hg38+rDNA reference end being mapped to the rDNA transcriptional unit, distribution of the origin of those reads across Hg38 contigs on the UKB prealigned files. The regions within grey boxes contain >98% of the reads.

**Figure S4.**
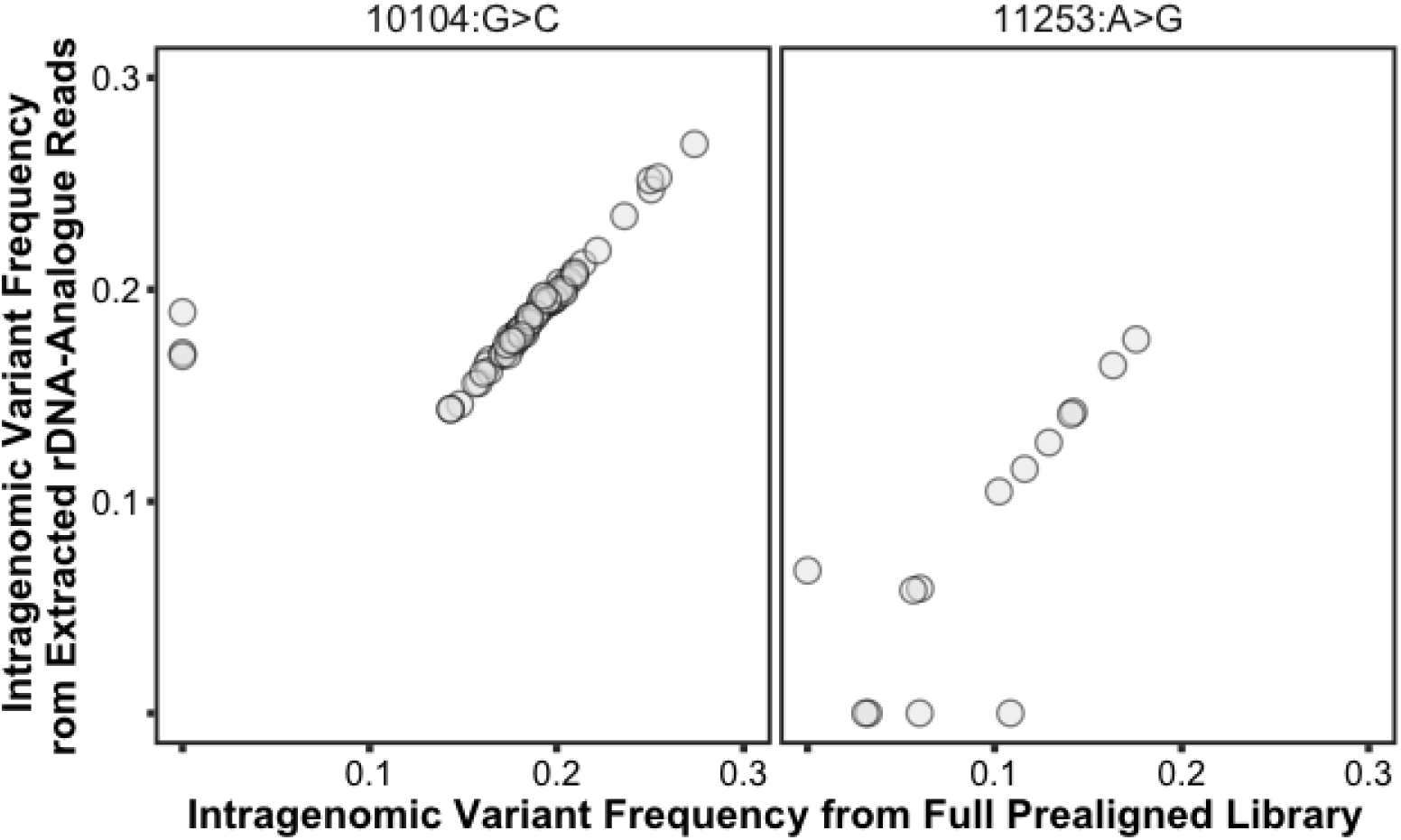
Comparison between the intragenomic frequencies in UKB MZ twin pairs obtained from full library realignments and prior extraction of reads from rDNA analogues, for two variants with R < 0.8 in such comparison.

**Figure S5.**
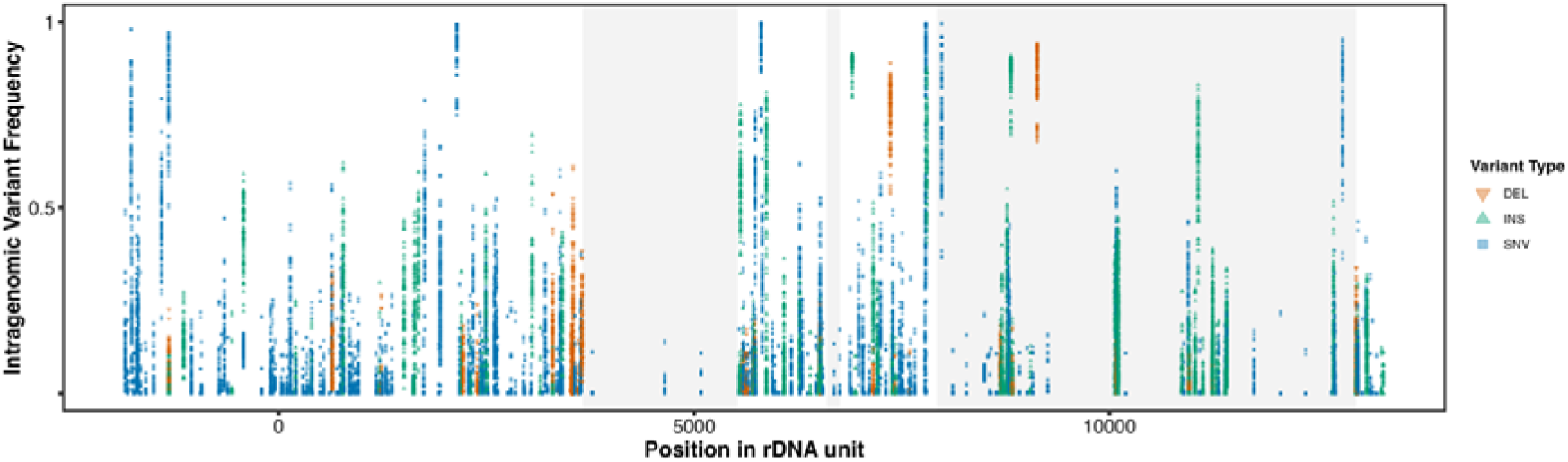
Distribution of intragenomic variant frequencies across the rDNA transcriptional unit for variants selected for further analysis.

**Figure S6.**
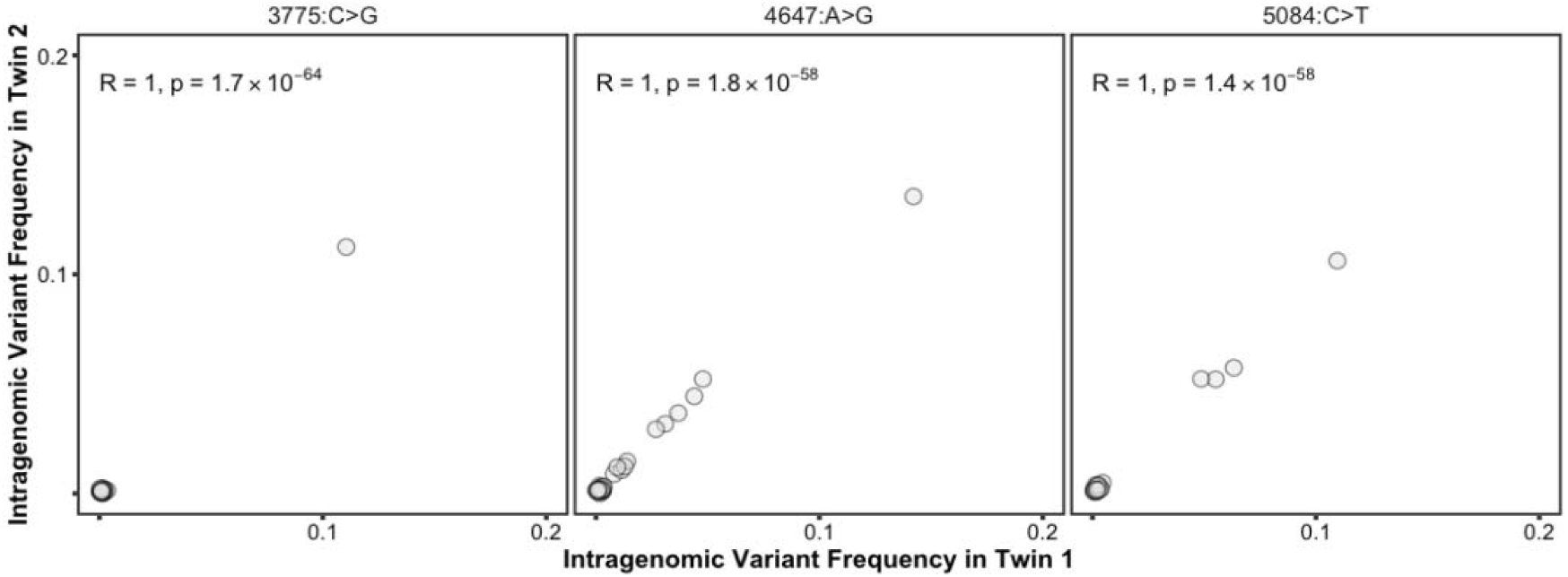
Comparison of estimated intragenomic variant frequencies in MZ twin pairs for selected variants from within the 18S.

**Figure S7.**
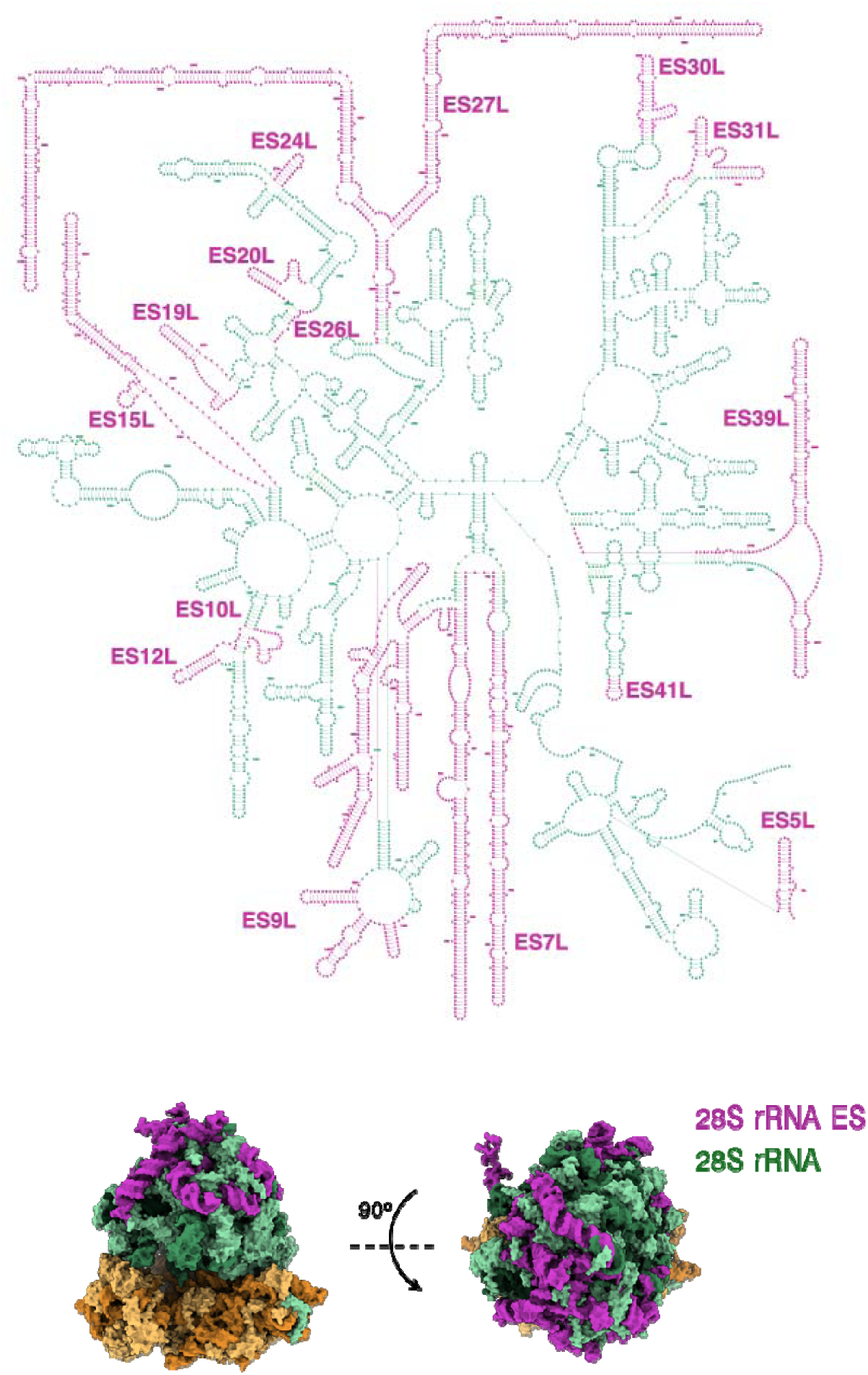
Modelling of the location of 28S expansion segments (ES) both within the 2D rRNA (Top) and the 3D ribosome (Bottom) structures. Ribosome models generated from PDB accession 8QOI^[55]^.

**Figure S8.**
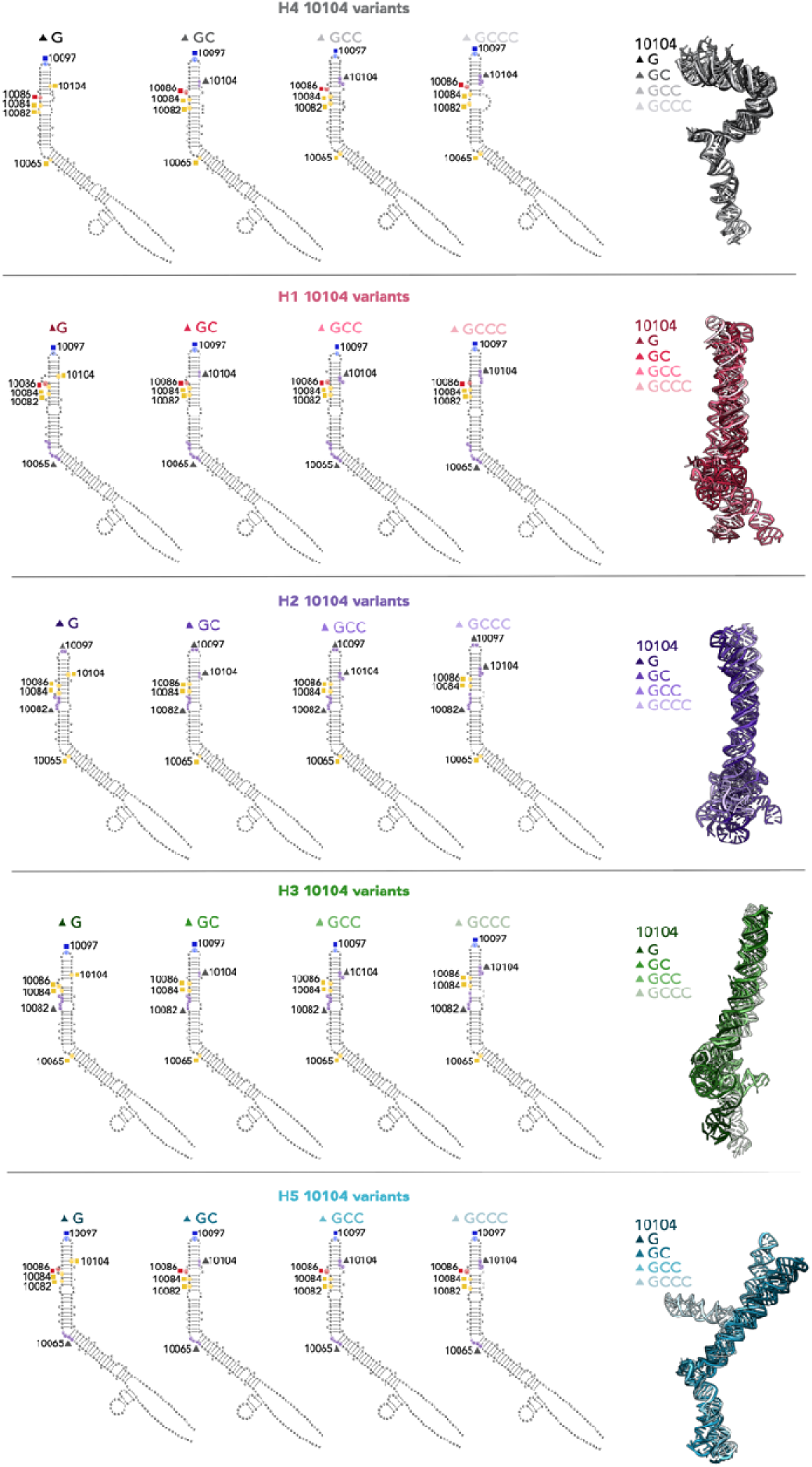
Predicted secondary and tertiary structures arising from the variants at position 10104, for ES15L haplotypes H1-5. For each haplotype and 10104 variant, the predicted 3D structures are aligned and given on the right. Symbols and colours indicate variants as in Fig. 3E.

**Figure S9.**
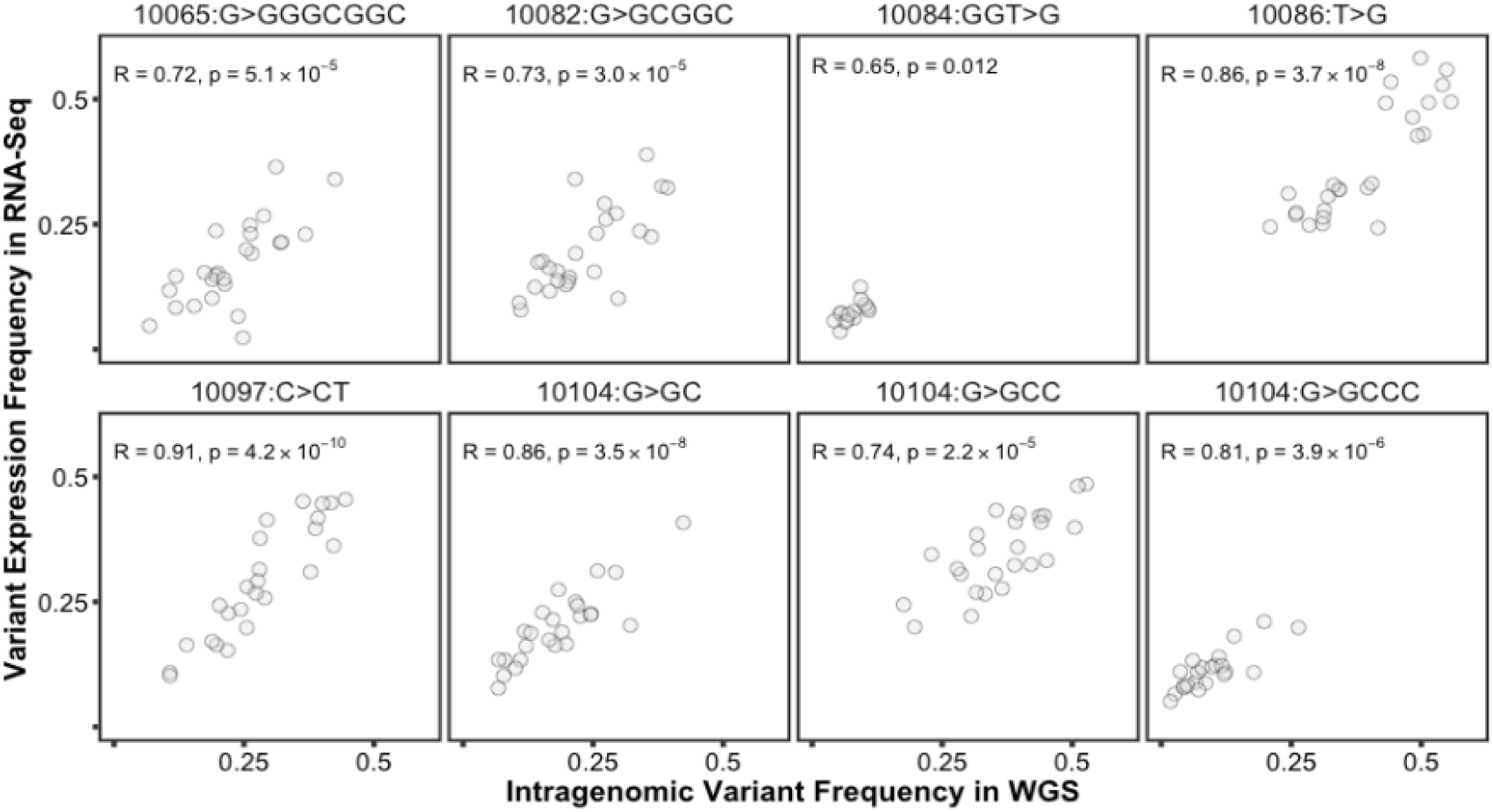
Comparison between variant frequencies obtained on DNA and RNA data from GBR participants of the 1000 Genomes Project, for UKB phenotype-associated variants in ES15L.

**Figure S10.**
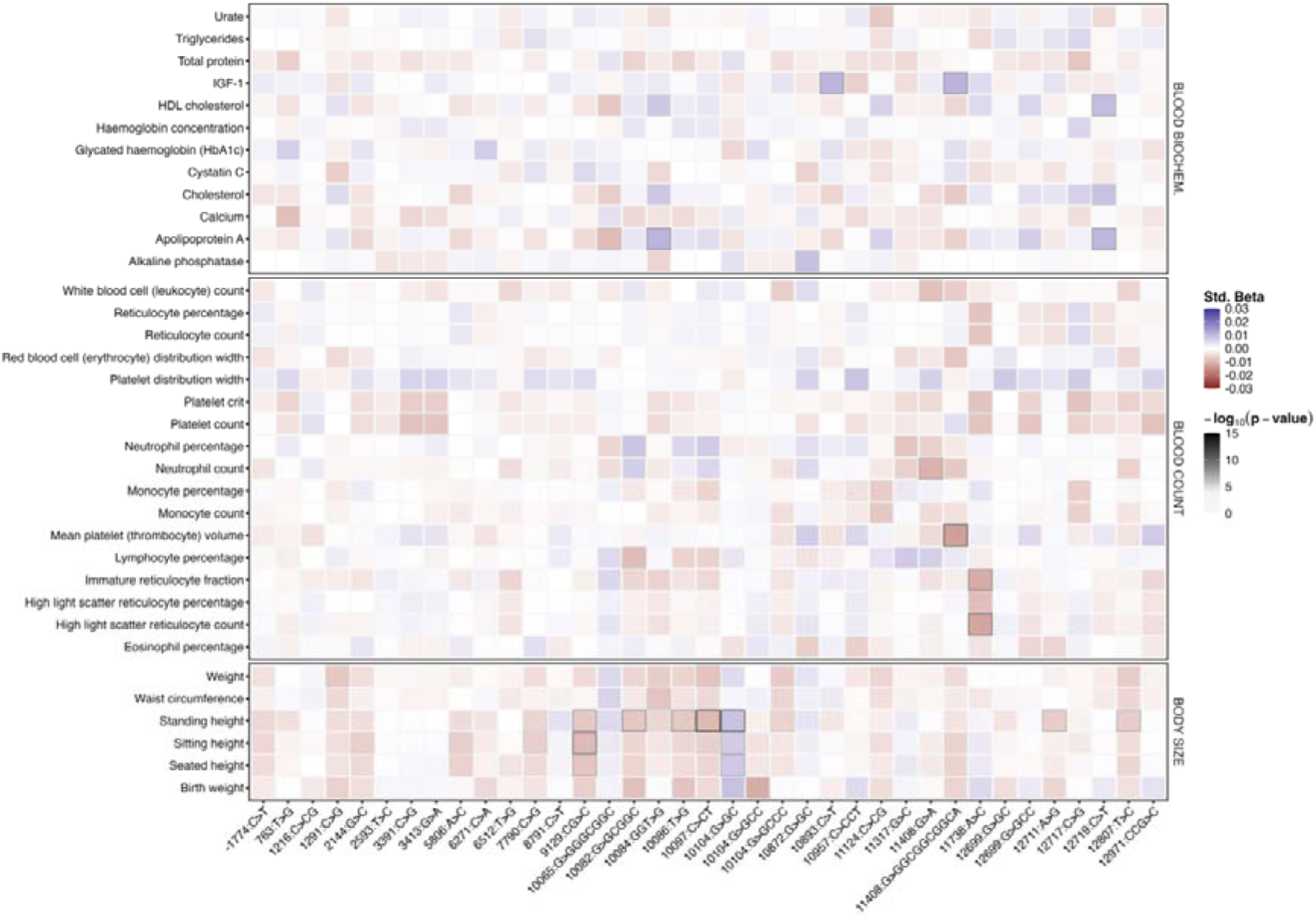
Heatmap of effect sizes obtained from regression models of variant frequency for combinations of variants and phenotypes with per-variant associations included in Fig. 3B.

**Figure S11.**
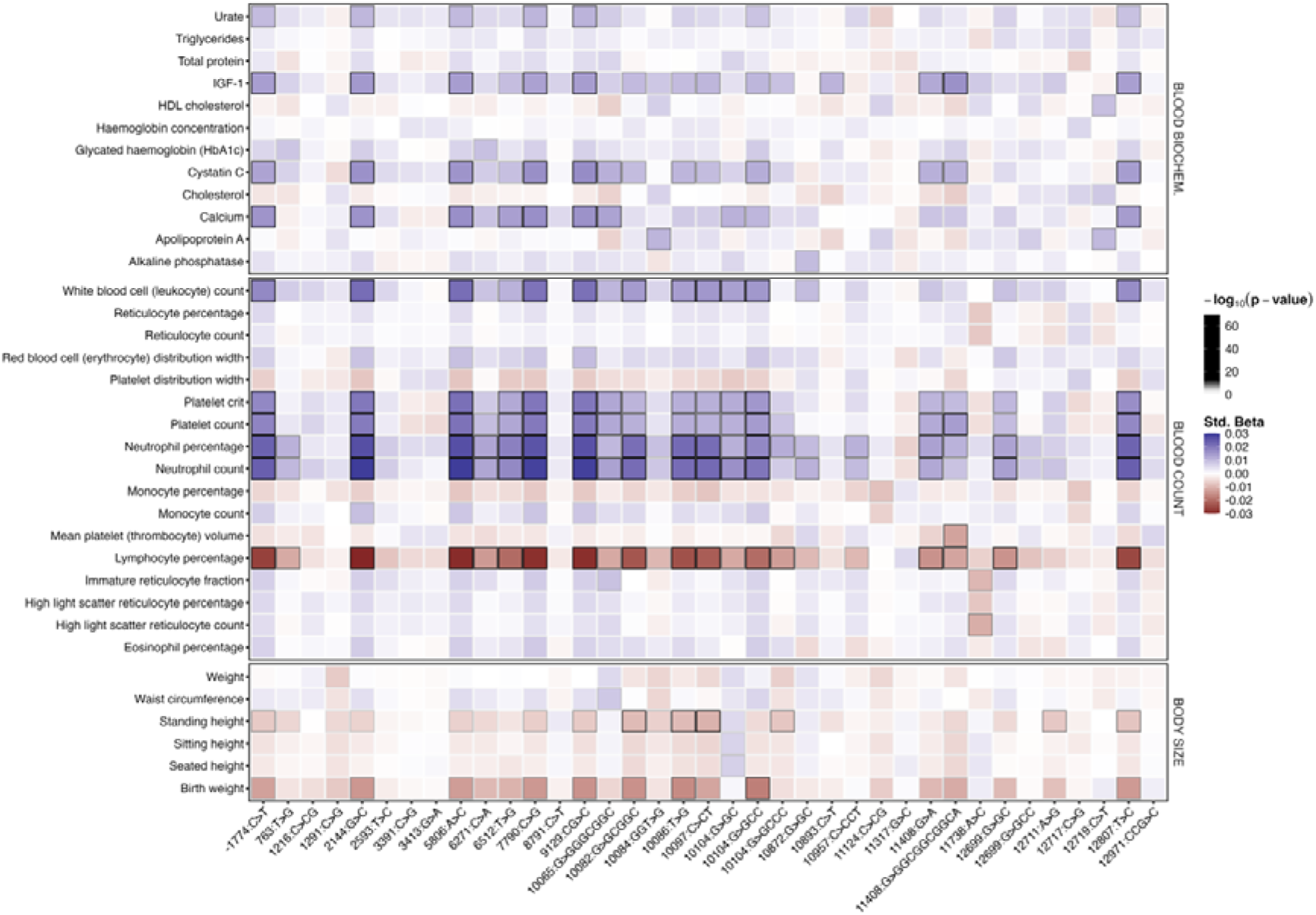
Heatmap of effect sizes obtained from regression models of allele-specific copy number for combinations of variants and phenotypes with per-variant associations included in Fig. 3B.

**Figure S12.**
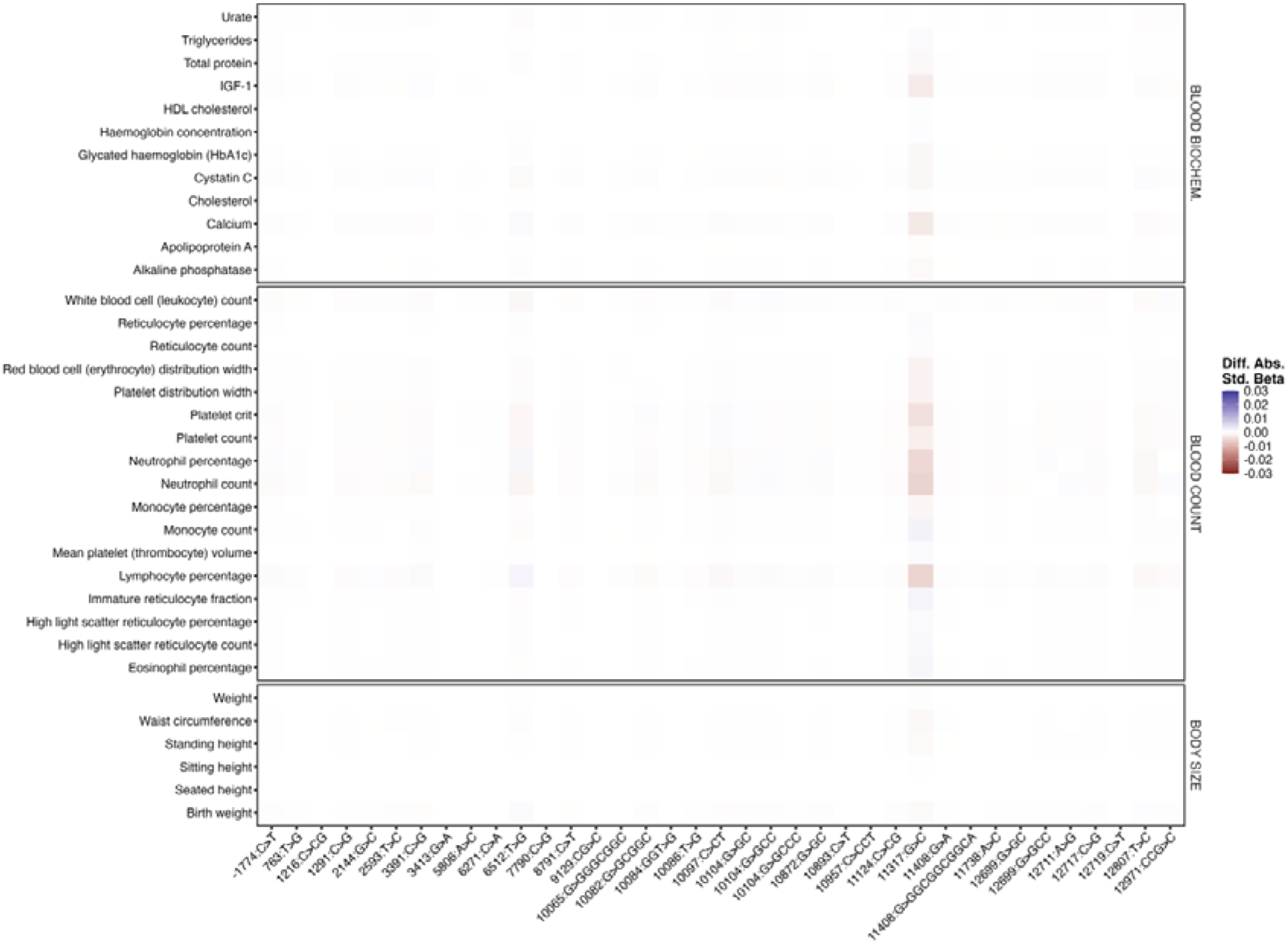
Heatmap of differences in absolute effect sizes obtained from regression models of variant frequency with and without total rDNA CN as covariate, for combinations of variants and phenotypes with per-variant associations included in Fig. 3B.

**Figure S13.**
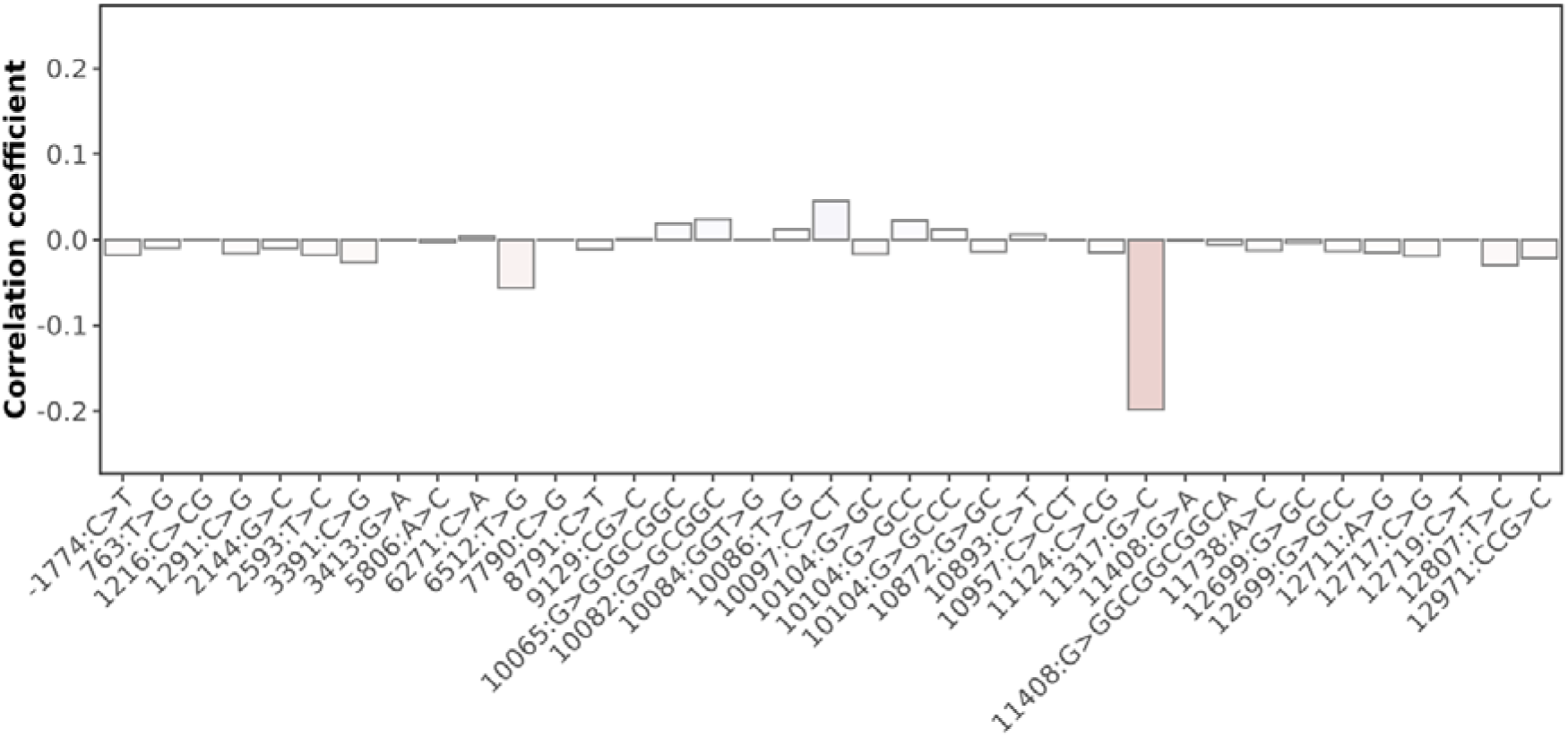
For the variants included in Fig. 3B, Pearson’s R estimates for the correlation between the corresponding intragenomic variant frequency and the total rDNA CN for unrelated UKB white British participants.

**Figure S14.**
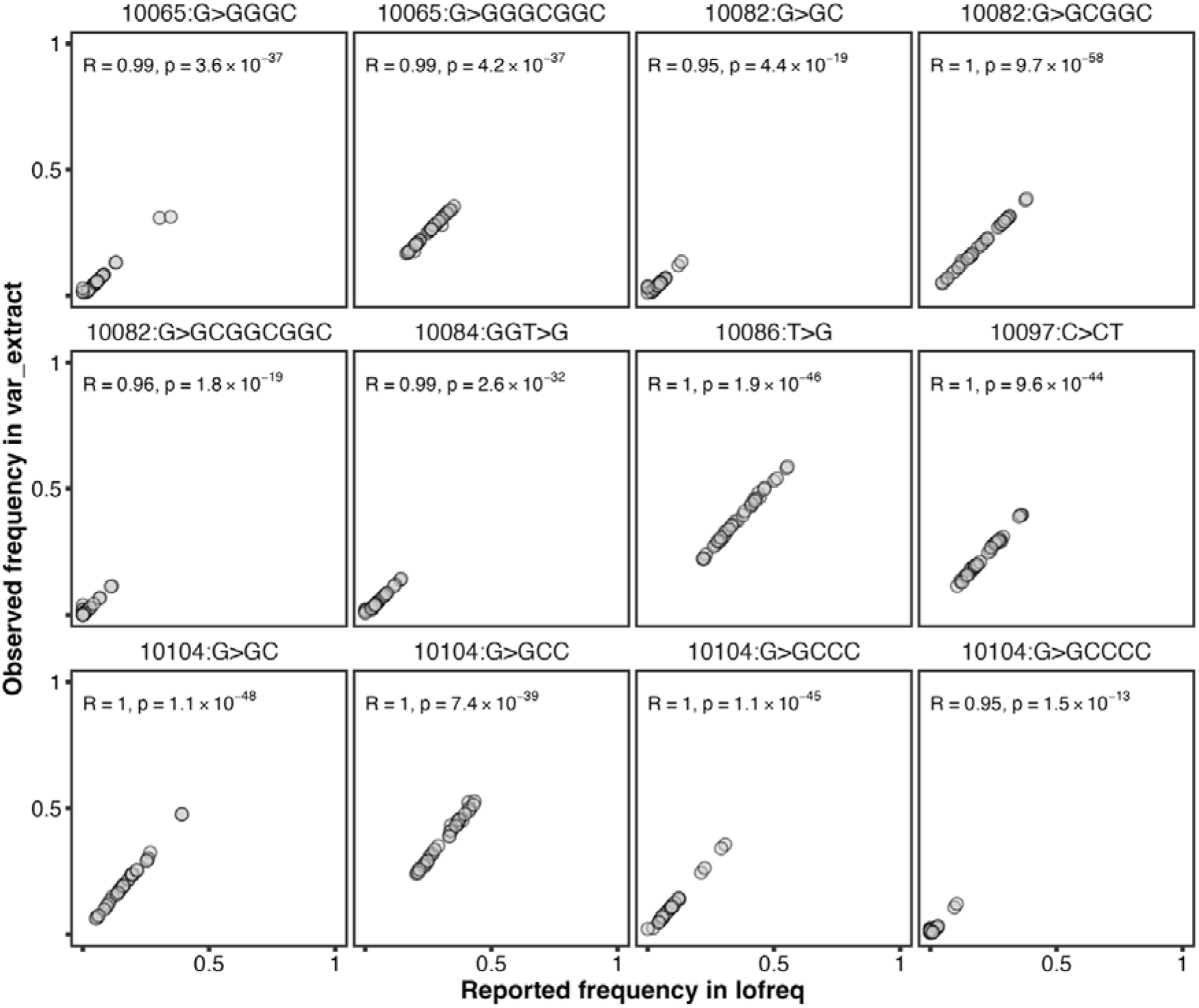
Comparison between ES15L variant frequencies obtained from `lofreq` and those derived from the per-read variant calls from the in-house `var_extract` script on MZ twin pairs included in the second deCODE WGS release.

## References

1. Parks, M. M., Kurylo, C. M., Dass, R. A., Bojmar, L., Lyden, D., Vincent, C. T., & Blanchard, S. C. (2018). Variant ribosomal RNA alleles are conserved and exhibit tissue-specific expression. Science Advances, 4(2), eaao0665. 10.1126/sciadv.aao0665

2. Hall, A. N., Morton, E., & Queitsch, C. (2022). First discovered, long out of sight, finally visible: Ribosomal DNA. Trends in Genetics, 38(6), 587–597. 10.1016/j.tig.2022.02.005

3. Fan, W., Eklund, E., Sherman, R. M., Liu, H., Pitts, S., Ford, B., Rajeshkumar, N. V., & Laiho, M. (2022). Widespread genetic heterogeneity of human ribosomal RNA genes. RNA, 28(4), 478–492. 10.1261/rna.078925.121

4. Rothschild, D., Susanto, T. T., Sui, X., Spence, J. P., Rangan, R., Genuth, N. R., Sinnott-Armstrong, N., Wang, X., Pritchard, J. K., & Barna, M. (2024). Diversity of ribosomes at the level of rRNA variation associated with human health and disease. Cell Genomics, 4(9). 10.1016/j.xgen.2024.100629

5. Wang, W., Zhang, X., Garcia, S., Leitch, A. R., & Kovařík, A. (2023). Intragenomic rDNA variation—The product of concerted evolution, mutation, or something in between? Heredity, 131(3), 179–188. 10.1038/s41437-023-00634-5

6. Rodriguez-Algarra, F., Evans, D. M., & Rakyan, V. K. (2024). Ribosomal DNA copy number variation associates with hematological profiles and renal function in the UK Biobank. Cell Genomics, 4(6). 10.1016/j.xgen.2024.100562

7. Rodriguez-Algarra, F., Whittaker, E., del Castillo del Rio, S., & Rakyan, V. K. (2025). Assessing Human Ribosomal DNA Variation and Its Association With Phenotypic Outcomes. BioEssays, n/a(n/a), e202400232. 10.1002/bies.202400232

8. Wang, M., & Lemos, B. (2017). Ribosomal DNA copy number amplification and loss in human cancers is linked to tumor genetic context, nucleolus activity, and proliferation. PLOS Genetics, 13(9), e1006994. 10.1371/journal.pgen.1006994

9. Xu, B., Li, H., Perry, J. M., Singh, V. P., Unruh, J., Yu, Z., Zakari, M., McDowell, W., Li, L., & Gerton, J. L. (2017). Ribosomal DNA copy number loss and sequence variation in cancer. PLOS Genetics, 13(6), e1006771. 10.1371/journal.pgen.1006771

10. Zafiropoulos, A., Tsentelierou, E., Linardakis, M., Kafatos, A., & Spandidos, D. A. (2005). Preferential loss of 5S and 28S rDNA genes in human adipose tissue during ageing. The International Journal of Biochemistry & Cell Biology, 37(2), 409–415. 10.1016/j.biocel.2004.07.007

11. Feng, L., Du, J., Yao, C., Jiang, Z., Li, T., Zhang, Q., Guo, X., Yu, M., Xia, H., Shi, L., Jia, J., Tong, Y., Ju, L., Liu, J., Lou, J., & Lemos, B. (2020). Ribosomal DNA copy number is associated with P53 status and levels of heavy metals in gastrectomy specimens from gastric cancer patients. Environment International, 138, 105593. 10.1016/j.envint.2020.105593

12. Lou, J., Yu, S., Feng, L., Guo, X., Wang, M., Branco, A. T., Li, T., & Lemos, B. (2021). Environmentally induced ribosomal DNA (rDNA) instability in human cells and populations exposed to hexavalent chromium [Cr (VI)]. Environment International, 153, 106525. 10.1016/j.envint.2021.106525

13. Law, P. P., Mikheeva, L. A., Rodriguez-Algarra, F., Asenius, F., Gregori, M., Seaborne, R. A. E., Yildizoglu, S., Miller, J. R. C., Tummala, H., Mesnage, R., Antoniou, M. N., Li, W., Tan, Q., Hillman, S. L., Rakyan, V. K., Williams, D. J., & Holland, M. L. (2024). Ribosomal DNA copy number is associated with body mass in humans and other mammals. Nature Communications, 15(1), 5006. 10.1038/s41467-024-49397-5

14. Xue, S., & Barna, M. (2012). Specialized ribosomes: A new frontier in gene regulation and organismal biology. Nature Reviews Molecular Cell Biology, 13(6), Article 6. 10.1038/nrm3359

15. Gay, D. M., Lund, A. H., & Jansson, M. D. (2022). Translational control through ribosome heterogeneity and functional specialization. Trends in Biochemical Sciences, 47(1), 66–81. 10.1016/j.tibs.2021.07.001

16. Miller, S. C., MacDonald, C. C., Kellogg, M. K., Karamysheva, Z. N., & Karamyshev, A. L. (2023). Specialized Ribosomes in Health and Disease. International Journal of Molecular Sciences, 24(7), 6334. 10.3390/ijms24076334

17. Kurylo, C. M., Parks, M. M., Juette, M. F., Zinshteyn, B., Altman, R. B., Thibado, J. K., Vincent, C. T., & Blanchard, S. C. (2018). Endogenous rRNA Sequence Variation Can Regulate Stress Response Gene Expression and Phenotype. Cell Reports, 25(1), 236–248.e6. 10.1016/j.celrep.2018.08.093

18. Song, W., Joo, M., Yeom, J.-H., Shin, E., Lee, M., Choi, H.-K., Hwang, J., Kim, Y.-I., Seo, R., Lee, J. E., Moore, C. J., Kim, Y.-H., Eyun, S., Hahn, Y., Bae, J., & Lee, K. (2019). Divergent rRNAs as regulators of gene expression at the ribosome level. Nature Microbiology, 4(3), 515–526. 10.1038/s41564-018-0341-1

19. Bycroft, C., Freeman, C., Petkova, D., Band, G., Elliott, L. T., Sharp, K., Motyer, A., Vukcevic, D., Delaneau, O., O’Connell, J., Cortes, A., Welsh, S., Young, A., Effingham, M., McVean, G., Leslie, S., Allen, N., Donnelly, P., & Marchini, J. (2018). The UK Biobank resource with deep phenotyping and genomic data. Nature, 562(7726), 203–209. 10.1038/s41586-018-0579-z

20. Halldorsson, B. V., Eggertsson, H. P., Moore, K. H. S., Hauswedell, H., Eiriksson, O., Ulfarsson, M. O., Palsson, G., Hardarson, M. T., Oddsson, A., Jensson, B. O., Kristmundsdottir, S., Sigurpalsdottir, B. D., Stefansson, O. A., Beyter, D., Holley, G., Tragante, V., Gylfason, A., Olason, P. I., Zink, F., … Stefansson, K. (2022). The sequences of 150,119 genomes in the UK Biobank. Nature, 607(7920), 732–740. 10.1038/s41586-022-04965-x

21. Li, S., Carss, K. J., Halldorsson, B. V., Cortes, A., & Consortium, U. B. W.-G. S. (2023). Whole-genome sequencing of half-a-million UK Biobank participants (p. 2023.12.06.23299426). medRxiv. 10.1101/2023.12.06.23299426

22. Zentner, G. E., Saiakhova, A., Manaenkov, P., Adams, M. D., & Scacheri, P. C. (2011). Integrative genomic analysis of human ribosomal DNA. Nucleic Acids Research, 39(12), 4949–4960. 10.1093/nar/gkq1326

23. Rodriguez-Algarra, F., Seaborne, R. A. E., Danson, A. F., Yildizoglu, S., Yoshikawa, H., Law, P. P., Ahmad, Z., Maudsley, V. A., Brew, A., Holmes, N., Ochôa, M., Hodgkinson, A., Marzi, S. J., Pradeepa, M. M., Loose, M., Holland, M. L., & Rakyan, V. K. (2022). Genetic variation at mouse and human ribosomal DNA influences associated epigenetic states. Genome Biology, 23(1), 54. 10.1186/s13059-022-02617-x

24. George, S. S., Pimkin, M., & Paralkar, V. R. (2023). Construction and validation of customized genomes for human and mouse ribosomal DNA mapping. Journal of Biological Chemistry, 299(6). 10.1016/j.jbc.2023.104766

25. Kasselimi, E., Pefani, D.-E., Taraviras, S., & Lygerou, Z. (2022). Ribosomal DNA and the nucleolus at the heart of aging. Trends in Biochemical Sciences, 47(4), 328–341. 10.1016/j.tibs.2021.12.007

26. Peterson, C. R. D., Cryar, J. R., & Gaubatz, J. W. (1984). Constancy of ribosomal RNA genes during aging of mouse heart cells and during serial passage of WI-38 cells. Archives of Gerontology and Geriatrics, 3(2), 115–125. 10.1016/0167-4943(84)90004-9

27. Malinovskaya, E. M., Ershova, E. S., Golimbet, V. E., Porokhovnik, L. N., Lyapunova, N. A., Kutsev, S. I., Veiko, N. N., & Kostyuk, S. V. (2018). Copy Number of Human Ribosomal Genes With Aging: Unchanged Mean, but Narrowed Range and Decreased Variance in Elderly Group. Frontiers in Genetics, 9. 10.3389/fgene.2018.00306

28. Qu, L.-H., Nicoloso, M., & Bachellerie, J.-P. (1991). A sequence dimorphism in a conserved domain of human 28S rRNA. Uneven distribution of variant genes among individuals. Differential expression in HeLa cells. Nucleic Acids Research, 19(5), 1015–1019. 10.1093/nar/19.5.1015

29. Wilm, A., Aw, P. P. K., Bertrand, D., Yeo, G. H. T., Ong, S. H., Wong, C. H., Khor, C. C., Petric, R., Hibberd, M. L., & Nagarajan, N. (2012). LoFreq: A sequence-quality aware, ultra-sensitive variant caller for uncovering cell-population heterogeneity from high-throughput sequencing datasets. Nucleic Acids Research, 40(22), 11189–11201. 10.1093/nar/gks918

30. Millard, L. A., Davies, N. M., Gaunt, T. R., Davey Smith, G., & Tilling, K. (2018). Software Application Profile: PHESANT: a tool for performing automated phenome scans in UK Biobank. International Journal of Epidemiology, 47(1), 29–35. 10.1093/ije/dyx204

31. Gonzalez, I. L., Sylvester, J. E., & Schmickel, R. D. (1988). Human 28S ribosomal RNA sequence heterogeneity. Nucleic Acids Research, 16(21), 10213–10224. 10.1093/nar/16.21.10213

32. Ramesh, M., & Woolford, J. L. (2016). Eukaryote-specific rRNA expansion segments function in ribosome biogenesis. RNA, 22(8), 1153–1162. 10.1261/rna.056705.116

33. Fujii, K., Susanto, T. T., Saurabh, S., & Barna, M. (2018). Decoding the Function of Expansion Segments in Ribosomes. Molecular Cell, 72(6), 1013–1020.e6. 10.1016/j.molcel.2018.11.023

34. Parker, M. S., Balasubramaniam, A., Sallee, F. R., & Parker, S. L. (2018). The Expansion Segments of 28S Ribosomal RNA Extensively Match Human Messenger RNAs. Frontiers in Genetics, 9. 10.3389/fgene.2018.00066

35. Genuth, N. R., & Barna, M. (2018). Heterogeneity and specialized functions of translation machinery: From genes to organisms. Nature Reviews Genetics, 19(7), 431–452. 10.1038/s41576-018-0008-z

36. Sauert, M., Temmel, H., & Moll, I. (2015). Heterogeneity of the translational machinery: Variations on a common theme. Biochimie, 114, 39–47. 10.1016/j.biochi.2014.12.011

37. Cui, L., Zheng, J., Lin, Y., Lin, P., Lu, Y., Zheng, Y., Guo, B., & Zhao, X. (2024). Decoding the ribosome’s hidden language: rRNA modifications as key players in cancer dynamics and targeted therapies. Clinical and Translational Medicine, 14(5), e1705. 10.1002/ctm2.1705

38. Wickham, H., Averick, M., Bryan, J., Chang, W., McGowan, L. D., François, R., Grolemund, G., Hayes, A., Henry, L., Hester, J., Kuhn, M., Pedersen, T. L., Miller, E., Bache, S. M., Müller, K., Ooms, J., Robinson, D., Seidel, D. P., Spinu, V., … Yutani, H. (2019). Welcome to the Tidyverse. Journal of Open Source Software, 4(43), 1686. 10.21105/joss.01686

39. Li, H., Handsaker, B., Wysoker, A., Fennell, T., Ruan, J., Homer, N., Marth, G., Abecasis, G., Durbin, R., & 1000 Genome Project Data Processing Subgroup. (2009). The Sequence Alignment/Map format and SAMtools. Bioinformatics, 25(16), 2078–2079. 10.1093/bioinformatics/btp352

40. Holtgrewe, M. (2010). Mason – A Read Simulator for Second Generation Sequencing Data. Technical Report FU Berlin. https://publications.imp.fu-berlin.de/962/

41. Langmead, B., & Salzberg, S. L. (2012). Fast gapped-read alignment with Bowtie 2. Nature Methods, 9(4), 357–359. 10.1038/nmeth.1923

42. Quinlan, A. R., & Hall, I. M. (2010). BEDTools: A flexible suite of utilities for comparing genomic features. Bioinformatics, 26(6), 841–842. 10.1093/bioinformatics/btq033

43. Bushnell, B. (2022). BBMap (Version 38.95) [Computer software]. https://sourceforge.net/projects/bbmap/

44. Krueger, F. (2019). TrimGalore (Version 0.6.5) [Computer software]. https://github.com/FelixKrueger/TrimGalore

45. Taylor, D. J., Chhetri, S. B., Tassia, M. G., Biddanda, A., Yan, S. M., Wojcik, G. L., Battle, A., & McCoy, R. C. (2024). Sources of gene expression variation in a globally diverse human cohort. Nature, 632(8023), 122–130. 10.1038/s41586-024-07708-2

46. Gustafson, J. A., Gibson, S. B., Damaraju, N., Zalusky, M. P. G., Hoekzema, K., Twesigomwe, D., Yang, L., Snead, A. A., Richmond, P. A., Coster, W. D., Olson, N. D., Guarracino, A., Li, Q., Miller, A. L., Goffena, J., Anderson, Z. B., Storz, S. H. R., Ward, S. A., Sinha, M., … Miller, D. E. (2024). High-coverage nanopore sequencing of samples from the 1000 Genomes Project to build a comprehensive catalog of human genetic variation. Genome Research, 34(11), 2061–2073. 10.1101/gr.279273.124

47. Oxford Nanopore Technologies. (2024). Pod5: A high performance file format for nanopore reads [Computer software]. https://github.com/nanoporetech/pod5-file-format

48. Li, H. (2018). Minimap2: Pairwise alignment for nucleotide sequences. Bioinformatics, 34(18), 3094–3100. 10.1093/bioinformatics/bty191

49. Li, H. (2021). New strategies to improve minimap2 alignment accuracy. Bioinformatics, 37(23), 4572–4574. 10.1093/bioinformatics/btab705

50. Oxford Nanopore Technologies. (2019). Megalodon [Computer software]. https://github.com/nanoporetech/megalodon

51. Sweeney, B. A., Hoksza, D., Nawrocki, E. P., Ribas, C. E., Madeira, F., Cannone, J. J., Gutell, R., Maddala, A., Meade, C. D., Williams, L. D., Petrov, A. S., Chan, P. P., Lowe, T. M., Finn, R. D., & Petrov, A. I. (2021). R2DT is a framework for predicting and visualising RNA secondary structure using templates. Nature Communications, 12(1), 3494. 10.1038/s41467-021-23555-5

52. Johnson, P. Z., & Simon, A. E. (2023). RNAcanvas: Interactive drawing and exploration of nucleic acid structures. Nucleic Acids Research, 51(W1), W501–W508. 10.1093/nar/gkad302

53. Wang, W., Feng, C., Han, R., Wang, Z., Ye, L., Du, Z., Wei, H., Zhang, F., Peng, Z., & Yang, J. (2023). trRosettaRNA: Automated prediction of RNA 3D structure with transformer network. Nature Communications, 14(1), 7266. 10.1038/s41467-023-42528-4

54. Pettersen, E. F., Goddard, T. D., Huang, C. C., Meng, E. C., Couch, G. S., Croll, T. I., Morris, J. H., & Ferrin, T. E. (2021). UCSF ChimeraX: Structure visualization for researchers, educators, and developers. Protein Science, 30(1), 70–82. 10.1002/pro.3943

55. Holvec, S., Barchet, C., Lechner, A., Fréchin, L., De Silva, S. N. T., Hazemann, I., Wolff, P., von Loeffelholz, O., & Klaholz, B. P. (2024). The structure of the human 80S ribosome at 1.9 Å resolution reveals the molecular role of chemical modifications and ions in RNA. Nature Structural & Molecular Biology, 31(8), 1251–1264. 10.1038/s41594-024-01274-x

